# Growth rate controls the sensitivity of gene regulatory circuits

**DOI:** 10.1101/2022.04.03.486858

**Authors:** Thomas Julou, Théo Gervais, Daan de Groot, Erik van Nimwegen

## Abstract

Unicellular organisms adapt to their changing environments by gene regulatory switches that sense chemical cues and induce specific target genes when the inducing signal is over a critical threshold. Using mathematical modeling we here show that, because growth rate sets the dilution rate of intracellular molecules, the sensitivity of gene regulatory switches automatically couples to growth rate, in a way that can be exploited by natural selection. We confirm the modeling predictions by experimentally demonstrating that, as nutrient quality is varied, the concentration of inducer required for activating the *lac* operon in *E. coli* increases quadratically with the population’s growth rate. Our theory further predicts that, when growth rate is instead modulated by translation inhibition, critical inducer levels are invariant, and we experimentally validate this prediction as well. Moreover, we establish that this growth-coupled sensitivity allows bacteria to implement concentration-dependent sugar preferences, in which a new carbon source is used only if its concentration is high enough to improve upon the current growth rate of the cells. Using microfluidics in combination with time-lapse microscopy, we validate experimentally that this strategy governs how mixtures of glucose and lactose are used in *E. coli* at single-cell level. Overall, growth-coupled sensitivity provides a general mechanism through which cells can ‘mute’ external signals in beneficial conditions when growth is fast, and become highly sensitive to alternative nutrients or stresses when growth is slow or arrested.

## Introduction

From bacteria to humans, most biological organisms sense signals in their environment and employ systems for processing this information to adapt their behavior. One challenge for such information processing systems is that optimal responses to environmental signals are often highly context-dependent. For example, whether it is worthwhile to pursue a particular nutrient when detecting it in the environment will crucially depend on whether other, better nutrients are also available. While the central nervous systems of multicellular eukaryotes obviously enable complex context-dependent responses, it is currently unclear to what extent bacteria are also capable of context-dependent responses to environmental stimuli, or how their relatively simple regulatory circuitry would implement such context-dependence.

A number of studies over the last decade has shown that bacteria obey several so called ‘growth laws’ which determine how major aspects of cell physiology and proteome allocation vary with the growth rate during exponential growth [1–4]. For example, it is well-known that as growth rates increases, the absolute sizes of cells increase and at high growth rates DNA becomes polyploid [5, 6]. However, as the rates of biomolecular reactions inside cells are generally a function of molecule concentrations, we here focus on the scaling of protein concentrations with growth rate.

Although the global effects of these growth laws on protein concentrations are broadly understood for constitutively expressed genes and target genes in elementary circuits such as those of constitutively expressed activators [7], to what extent growth rate affects the responses of the regulatory circuitry of cells to signals from their environment has so far not been explored. Here we use a combination of theoretical modeling and single-cell experiments with *E. coli* to show that growth rate is a key contextual parameter for the response of regulatory circuits and controls their sensitivity to external signals.

The origin of this *growth-coupled sensitivity* (GCS) lies in the effects of dilution by growth. For any intracellular molecule that is stable relative to the doubling time of the cell, the rate at which its concentration decays is dominated by dilution, and thus equal to the growth rate. Consequently, whenever the rate of production of an intracellular molecule does not also increase at least proportional to growth rate, its steady-state concentration will decrease with growth rate. For example, for molecules that are imported into the cell by membrane-bound transporters at a constant rate, and that are not actively degraded or metabolized, their steady-state concentrations will be inversely proportional to growth rate. Similarly, although transcription rates, mRNA decay rates and translation all scale in a non-trivial manner with growth rate, the protein concentrations of constitutively expressed genes have been shown to decrease approximately inversely with growth rate [2, 7]. For other categories of genes, the situation can be more complex. For example, for genes involved in carbon catabolism, it has been shown that their concentration decreases linearly with growth rate when growth rate is modulated by carbon source quality, but increases proportional to growth rate when it is modulated by translation inhibition [3].

The behavior of gene regulatory circuits is thus expected to be intrinsically coupled to changes in growth rate, since the concentrations of their protein players (e.g. transcription factors) as well as of signalling molecules either activating or repressing them are affected by dilution due to growth. To explore such growth-coupled effects on regulatory circuitry, we focus on regulatory switches in which a positive feedback loop is coupled to a signal. These regulatory switches typically switch from an uninduced (‘off’) to an induced (‘on’) state when the intracellular concentration of the activating signal surpasses a critical concentration. Such regulatory switches are involved in many biological functions including the lysis-lysogeny switches employed by phages [8], the regulatory circuitry involved in carbon source utilization [9–11], competence [12], sporulation [12], and virulence [13]. In addition, a large fraction of the two-component signaling systems used by prokaryotes involve positive feedback loops and behave as regulatory switches.

By exploring simple theoretical models, we show that GCS can be exploited by natural selection in a variety of ways. In the simplest scenarios, the result of increasing dilution rates is to decrease the sensitivity of regulatory switches so that faster growing cells are relatively insensitive, while slow growing or growth arrested cells become hyper-sensitive to signals in their environment. However, by tuning the parameters of the regulatory circuitry, this default behavior can be modified or fine-tuned in a number of ways as we will explore below using the *lac* system in *E. coli*, which is arguably the archetypical example of a regulatory switch.

## Results

We first investigated how growth rate might affect the functioning of regulatory switches using mathematical models. We follow standard approaches for modeling simple regulatory circuits using deterministic coupled ordinary differential equations in which gene regulation is modeled by assuming protein production rates are Hill functions of the concentrations of regulators, e.g. [14, 15]. However, in contrast to typical analyses of such models, we focus on the effects of dilution and explore how steady-state gene expression levels not only depend on extracellular concentrations of signaling molecules, but also on growth rate.

For a minimal bistable regulatory system consisting of an operon containing a transcription factor (TF) that activates its own expression, the parameter regime for which the system exhibits bistability has been shown to depend on growth rate [7]. We here show that this simple positive feedback loop behaves as a regulatory switch as a function of growth rate, i.e. even without any coupling to an external signal (Fig. 1A-C; SI 1.1). As growth rate is decreased, the operon goes from being stably switched off at high growth rates, to bistable at moderately slow growth, to being stably switched on at very slow growth. Moreover, the growth rates at which these transitions occur can be tuned by the basal expression level of the promoter (Fig. 1B-C). Thus, the natural coupling to growth rate due to dilution turns a simple positive feedback loop into a regulatory switch that senses the cell’s growth rate. Although we are not aware of examples of regulatory switches that are solely controlled by growth rate, it is conceivable that phages could use such systems to switch on their lytic cycle as the growth rate of their host bacteria drops below a critical value.

**Figure 1.**
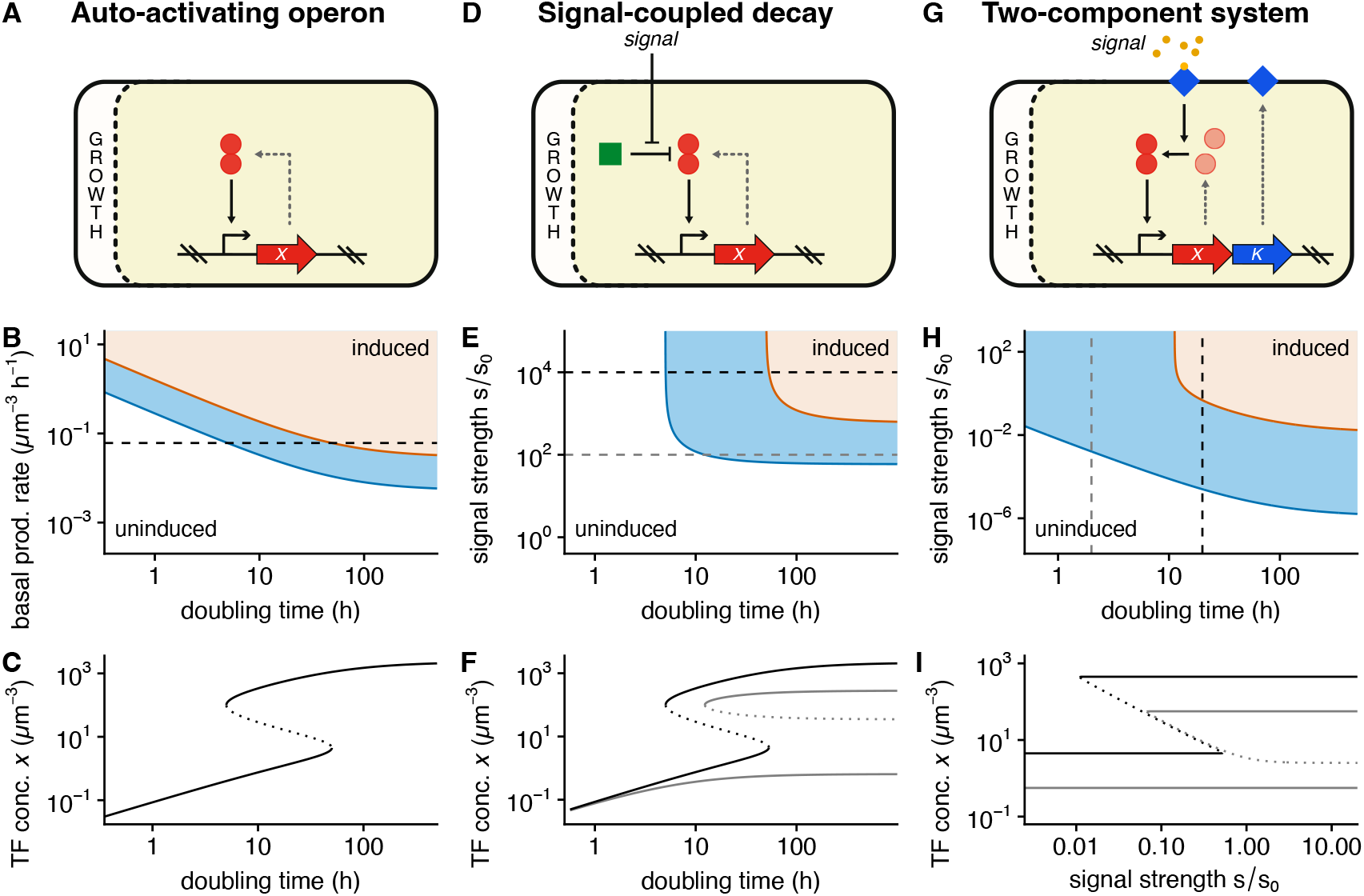
Theoretical analysis of gene regulatory switches exhibiting growth-coupled sensitivity due to growth rate dependent dilution. (A) A minimal regulatory switch circuit consisting of a TF (red circles) that positively regulates its own expression, coupled to growth rate through dilution of the TF. (B) Phase diagram of the auto-activating TF as a function of the doubling time and basal expression rate of the operon. Cells are uninduced in the white area of the plot, stably induced in the orange area, and bistable in the blue area. (C) Induction curves showing steady-state TF concentration *X* as a function of doubling time for a basal expression corresponding to the dotted line in panel B. The solid lines show the stable steady states and the dotted line the unstable steady state. Note that the system is bistable for doubling times between 5 and 50 hours. (D) A regulatory switch in which an auto-activating TF (red circles) is coupled to an external signal through active degradation by a protease (green square) whose activity is in turn repressed by an external signal. (E) Phase diagram for the regulatory switch with signal-coupled decay as a function of the doubling time and signal strength *s* relative to the signal strength *s*_0_ at which protease activity is at half-maximum. (F) Induction curves showing TF concentration as a function of doubling time at high signal strength (black curves and black dotted line in panel E) and intermediate signal strength (gray curves and gray dotted line in panel E). Note that at the intermediate signal strength, the system remains bistable for arbitrarily large doubling times. (G) Two-component system regulatory switch. Here the auto-activating operon contains a TF gene *X* (red circles) and a gene for a membrane-bound kinase *K* (blue diamonds). Kinase activity depends on an external signal and phosphorylation of the TF leads to its activation through dimerization. (H) Phase diagram of the two-component system as a function of doubling time and signal strength. Note that the minimal signal strength at which bistability sets in decreases approximately quadratically with doubling time. (I) Induction curves showing TF expression as a function of signal strength either at relatively fast growth (gray curves and grey dotted line in panel H) or at slow growth (black curves and black dotted line in panel H). The solid lines show stable steady states and the dotted lines unstable steady states. Note that at the faster growth rate the system remains bistable for arbitrarily high signal strengths. See SI for detailed parameter settings of each circuit.

In many known examples of regulatory switches, an operon activating its own expression is coupled to an external signal. For example, the regulatory circuits that implement competence and sporulation in *B. subtilis* have at their core an auto-activating TF that is coupled to an external signal through repression of a protease that determines the TF’s decay rate (Fig. 1D) [12]. The phase diagram of this type of circuit shows that it causes cells to only commit to sporulation/competence when both the external signal and the doubling time are over a critical value (Fig. 1E-F; SI 1.2). Thus, even though this regulatory circuit only interacts with a single signal, the coupling to growth rate makes the response of this system context-dependent, effectively integrating two signals. Switching can either occur by varying the signal strength at a given growth rate, or by varying the growth rate at a given signal strength. Moreover, the switching behavior as a function of one of these variables depends on the value of the other variable. For example, while at high signal strengths the system eventually stably switches on at sufficiently slow growth, at lower signal strength the system remains bistable even at growth arrest (Fig. 1F), illustrating how such circuits can be selected to function in one regime or the other. More generally, by tuning the parameters of the system, cells can implement stochastic responses in which only a subset of the population commits at intermediate growth rates or signal strengths. Indeed, it is well established that competence is only induced in a subset of cells but not in the whole population while sporulation occurs in the majority of cells [16].

The most common form of regulatory switches in bacteria involve two-component systems in which a TF positively regulates its own expression and is coupled to an external signal through phospho-relay by a membrane-bound kinase (Fig. 1G). In this simple example we assume the TF dimerizes upon being phosphorylated by the membrane-bound kinase, which in turn is activated by an external signal. In the parameter regime chosen for this example (see SI section 1.3 and Fig S1), the system switches from off to bistable, and then to stably on as a function of signal strength at low growth rates (Fig. 1H-I, black curves), but remains bistable for arbitrarily high signal strengths at high growth rate (Fig. 1H-I, gray curves). In addition, the threshold level of the signal at which the system becomes bistable increases roughly quadratically with growth rate (Fig. 1H; SI 1.3). That is, as doubling time increases by ten-fold, the threshold level of the signal decreases by hundred-fold, showing that the sensitivity of the regulatory circuit strongly decreases with growth rate. This quadratic scaling roughly results from the fact that the concentrations of both the kinase and TF scale inversely with growth rate and holds as long as growth rate is large compared to the decay rate of these molecules.

While these three simple examples only scratch the surface of the possible ways in which growth rate can affect the functioning of regulatory circuits, they illustrate that theoretical modeling predicts that the natural coupling to growth rate through dilution can profoundly affect the sensitivity of gene regulatory switches.

To investigate experimentally whether regulatory switches in bacteria indeed exhibit growth rate dependent sensitivity, we focused on the *lac* operon of *E. coli*. The *lac* operon consists of three genes which are involved in the metabolism of galactosides such as lactose. Its expression is repressed by the TF LacI and can be induced by lactose or artificial galactosides such as thio-methylgalactoside (TMG), which are not metabolized: when TMG is present, it inhibits DNA binding of the repressor LacI, leading to increased expression of the operon, including the transporter LacY, which increases transport of TMG into the cell, creating a positive feedback loop (Fig. 2A). Consequently, the *lac* operon can switch from an uninduced to an induced state when the TMG concentration exceeds a threshold value. We adapted a simple mathematical model of the *lac* operon [9] to model the effects of dilution (SI section 2 and Fig S2). We found that, in the simplest scenario where the rate of *lac* protein production at full induction is independent of growth rate, the model predicts that the critical external level of TMG decreases quadratically with the doubling times of the cells as long as this time is short relative to the half lifes of the molecules (Fig. 2B). Intuitively, this quadratic dependence results from the fact that, in this regime, the intracellular TMG concentration is set by the ratio of import and dilution and that the import rate is itself also inversely proportional to growth rate, because LacY expression is also a balance between production and dilution.

**Figure 2.**
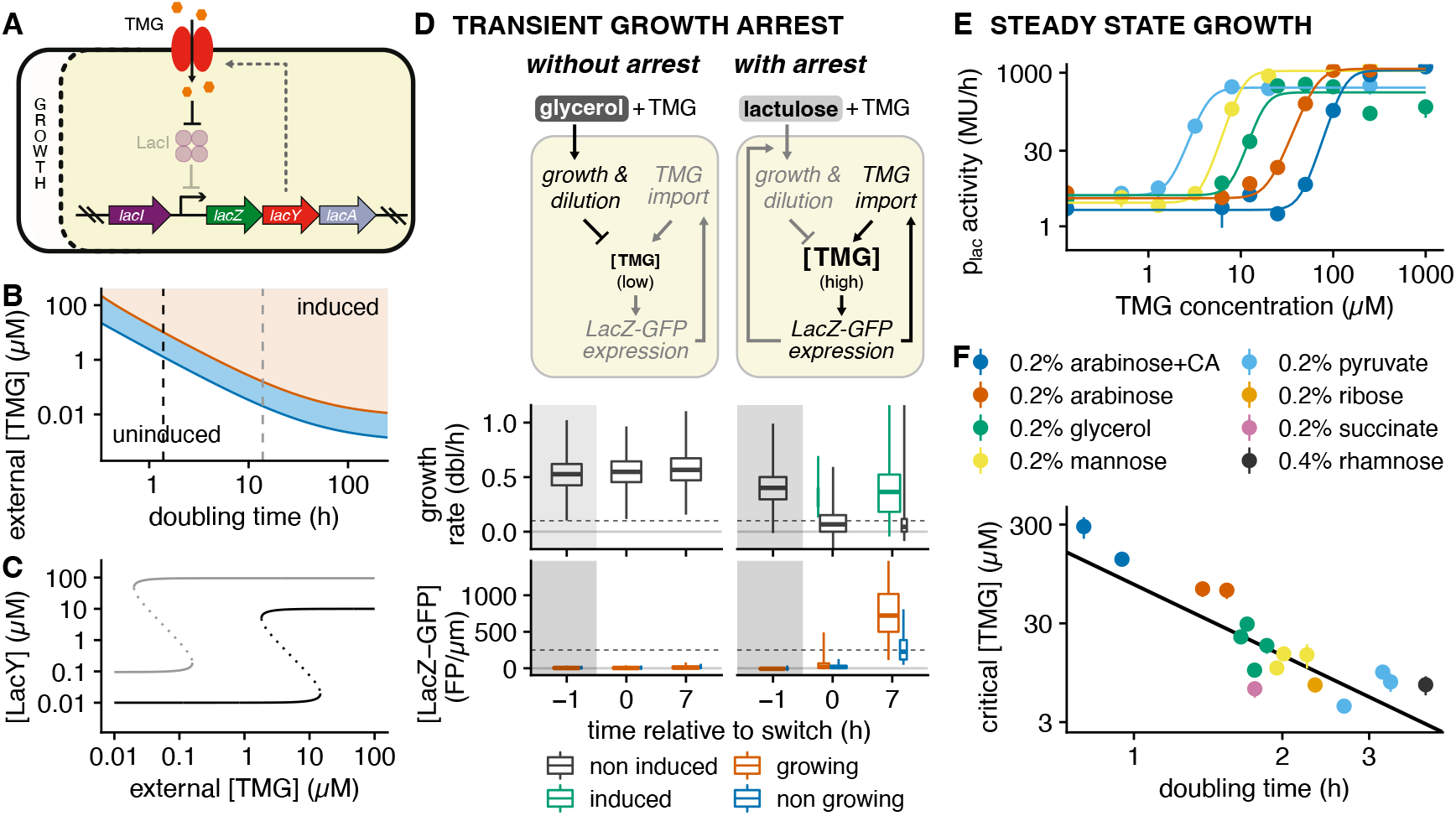
Growth-coupled sensitivity of the *lac* operon. **A**. Schematic of the *lac* operon’s positive feedback circuit coupled to the artificial inducer TMG. Note that the intracellular concentration of TMG is set by the balance between import by LacY and dilution due to growth, which leads to GCS. **B**. Theoretical phase diagram of *lac* operon expression as a function of doubling time and external inducer level (white: repressed, blue: bistable, orange: induced). The critical inducer level decreases quadratically with the doubling time until doubling time becomes comparable to the half life of TMG and the LacY proteins (here assumed to be 48 hours). **C**. Theoretically predicted induction curves of LacY expression as a function of inducer level at a 1 – 2 hour doubling time typical for growth in minimal media with lactose (black) and at a ten-fold higher doubling time (gray). See SI for detailed parameter values in B and C. **D**. Experimental validation that transient growth arrest increases the *lac* circuit’s sensitivity to TMG. Cells are initially grown in the Dual Input Mother Machine (DIMM) microfluidic device without TMG. The growth media are then switched to media that contain 20 µM of TMG and nutrients that either do not require *lac* operon expression (glycerol; left half), or nutrients that require *lac* operon expression but do not themselves induce *lac* operon expression (lactulose; right half). Note that in the latter cases cells enter growth arrest upon the switch in the media because the *lac* operon is initially repressed. The box-whisker plots show distributions of instantaneous growth rates (upper) and LacZ-GFP concentration (lower) immediately before, immediately after, and 7 h after the switch in the media (300 cells per time point). Note that cells were stratified by induction (upper) or growth rate (lower); see also Fig. S4. **E**. Induction curves of the *lac* operon as a function of TMG concentration in cells growing exponentially in M9 minimal media supplemented with different carbon sources, and where CRP activity is locked on (using a, 6.*cyaA*, 6.*cpdA* mutant supplemented with 1mM cAMP). Only one representative biological replicate is shown for each condition (all replicates are shown in Fig. S6); points and errors bars show the mean and standard error of three technical replicates; solid lines show fits to Hill functions; MU: Miller Units. **F**. The critical external concentration of TMG (estimated from fits as in E) as a function of doubling time shows an approximately quadratic decrease (exponent -2.4 ± 0.4) as predicted by our model (B).

Interestingly, this prediction reconciles observations in the literature that may have appeared contradictory. In particular, whereas Choi et al. [17] found critical levels of *lac* expression of several hundred molecules for cells growing in M9 media with glycerol and amino acids, our previous single-cell experiments showed that for growth arrested cells any nonzero *lac* expression is sufficient to induce the system when lactose is present [11]. Indeed, at parameter settings that match observations on the *lac* operon (SI section 2.3), the *lac* expression at the unstable steady state that separates the induced and uninduced states in the bistable region is predicted to be 0.1-0.2 µM (corresponding to a few hundred molecules) when cells are doubling every 1 - 2 hours, but at the same inducer level cells are guaranteed to induce at low growth rates or growth arrest (Fig. 2C).

One prediction of our theory is that, due to GCS, inducer concentrations exist that are too low to cause induction in growing cells, but that should cause induction when cells are growth arrested, even transiently. To test this prediction, we compared the responses of the *lac* to a given TMG concentration (20 µM TMG, see Fig. S3A) in 2 different media. Using microfluidics in combination with time-lapse microscopy [11, 18], we monitored growth and *lac* operon expression in single cells carrying a LacZ-GFP fusion at the native locus. Cells were initially grown in minimal media with glycerol and were then exposed either to the same media supplemented with 20 µM TMG, or to minimal media with lactulose and the same concentration of TMG (Fig. 2D). Notably, since lactulose requires LacZ to be metabolized, cells can only grow on lactulose if the *lac* operon is induced, but lactulose does not itself induce the *lac* operon [19]. Under the first change in media, cells continue to grow on glycerol and we observe that 20 µM of TMG is not sufficient to cause induction of the *lac* operon in any of the cells during the entire experiment (Fig. 2D). A very different behavior is observed for the second change in media. First, since the *lac* operon is repressed under growth on glycerol, the change to lactulose causes almost all cells to immediately go into growth arrest, but several hours later we observe that the large majority of cells have induced their *lac* operon and have recommenced growth (Fig. 2D, Fig. S3, and Fig. S4). Thus, the transient growth arrest led to the induction of the *lac* operon in the majority of cells, demonstrating experimentally that cells become more sensitive to inducers of the *lac* operon when they are growth-arrested.

We next set out to quantify how the critical external concentration of the inducer depends on growth rate: we grew batch cultures in media with different carbon sources to modulate growth rate and, in each of the growth media, measured *lac* expression as a function of TMG concentration using Miller assay (Fig. 2E and Fig. S5). Since we wanted to test the predictions of the GCS theory under the assumption of constant production at full induction, and since the *lac* operon is known to also be indirectly regulated by growth rate through CRP activity, we used a *cyaA cpdE* mutant strain in which CRP activity is kept constantly high by knocking out the synthesis and degradation of cAMP and by supplementing 1mM of extracellular cAMP [20, 21]. It has previously been observed that, when cAMP-CRP activity is kept constant, the expression of the *lac* operon at full induction indeed scales inversely proportional with growth rate [22].

Fitting Hill functions to the observed induction curves (Fig. 2E and Fig. S6), we estimated the critical external TMG concentration in each media. Remarkably, we found that the critical concentration at which the *lac* operon induces indeed decreases approximately quadratically with doubling time as predicted by the model (Fig. 2F and Fig. S7), with an almost 100-fold change in critical concentration between the fastest and slowest growth conditions. This confirms that the sensitivity of the *lac* operon to its inducers indeed depends strongly on the growth rate of the cells.

To check to what extent cAMP-CRP regulation affects the sensitivity of the *lac* operon, we performed analogous experiments with the wild-type strain. For this, we relied on fluorimetry using a strain carrying a LacZ-GFP fusion at the native locus (see SI section 9). We find that sensitivity decreases similarly with growth rate in the wild-type as in the mutant of cAMP-CRP regulation (Figs. S8 and S9). In addition, it is known that the activity of LacY transporters can be inhibited by other PTS sugar transporters ([23]) and this ‘inducer exclusion’ might also affect sensitivity of the *lac* system. However, using a mutant strain in which both cAMP-CRP activity is fixed and the crr gene is knocked out, we find that there is no systematic effect of inducer exclusion on either induction threshold or growth rate, and that growth rate remains the key determinant of the induction threshold. Finally, although it is impossible to fully rule out other growth rate dependent mechanisms, the match between the experiments and our simple model supports that this growth rate dependence is mainly mediated through the effects of dilution (see SI section 2.4).

While the foregoing experiments establish that regulatory switches such as the *lac* operon exhibit GCS, it is unclear to what extent evolution has exploited this GCS to tune regulatory responses in an adaptive manner. For example, the current predominant view is that *E. coli* has a fixed hierarchy of preferences for carbon sources, so that when a mixture of such carbon sources is available, cells first exhaust the preferred carbon source before consuming the less preferred one [10, 24]. Indeed, when grown in batch on a mixture of glucose and lactose, *E. coli* cells first consume glucose before starting to consume lactose. However, such a fixed hierarchy cannot always be optimal. For example, since it is well known that the growth rate that can be achieved on a given carbon source is a hyperbolic ‘Monod’ function of its concentration [25], the growth rates that can be attained on different carbon sources crucially depend on their concentrations, so that it cannot be optimal to always prefer a given carbon source over another independent of concentration. It is thus long been clear that the optimal choice of carbon source should be concentration-dependent, as has been observed by many others before us, e.g. [26, 27].

Ideally, cells should opt for whatever carbon source maximizes growth rate in a concentration-dependent manner. That is, if cells are growing on carbon source *A* at rate λ_*A*_, and an alternative carbon source *B* appears at concentration *c*_*B*_, then cells should ideally only switch to consuming *B* if the growth rate λ_*B*_ that they can attain on *B* is larger than the current growth rate λ_*A*_. Since the growth rate λ_*B*_ increases with concentration *c*_*B*_, the minimal concentration *c*_*B*_(λ_*A*_) that is required to ensure λ_*B*_ *≥*λ_*A*_ is an increasing function of the current growth rate λ_*A*_, and given by the inverse of the hyperbolic Monod function for carbon source *B*.

As we have seen above, GCS does naturally cause the critical concentration for inducing the regulatory switch to increase with growth rate, and this raises the intriguing question whether GCS might be tuned so as to achieve such optimal carbon source switching. That is, to ensure that the critical induction concentration as a function of growth rate for a given carbon source matches exactly the inverse of the Monod function for that carbon source. As detailed in section 3 of the SI, we find that this is indeed possible and is realized when the expression at full induction *y*_*h*_(λ) of the operon for the corresponding carbon source decreases linearly with growth rate λ as follows

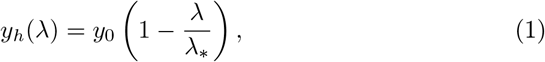

which should hold when growth rate is modulated by nutrient quality. In particular, when *y*_*h*_(λ) follows this form, it is possible for the critical external nutrient concentration to precisely track the inverse of the Monod curve provided that the parameters *y*_0_ and λ_*_ equal particular ratios of the kinetic parameters of the *lac* system (see equations S53 and S54 in SI section 3). The resulting optimal phase diagram is shown in Fig. 3A. Moreover, this behavior is also consistent with the relatively complex dynamics of LacZ converting lactose into the natural inducer allolactose, and its hydrolysis of both lactose and allolactose (SI section 4 and Fig. S10).

**Figure 3.**
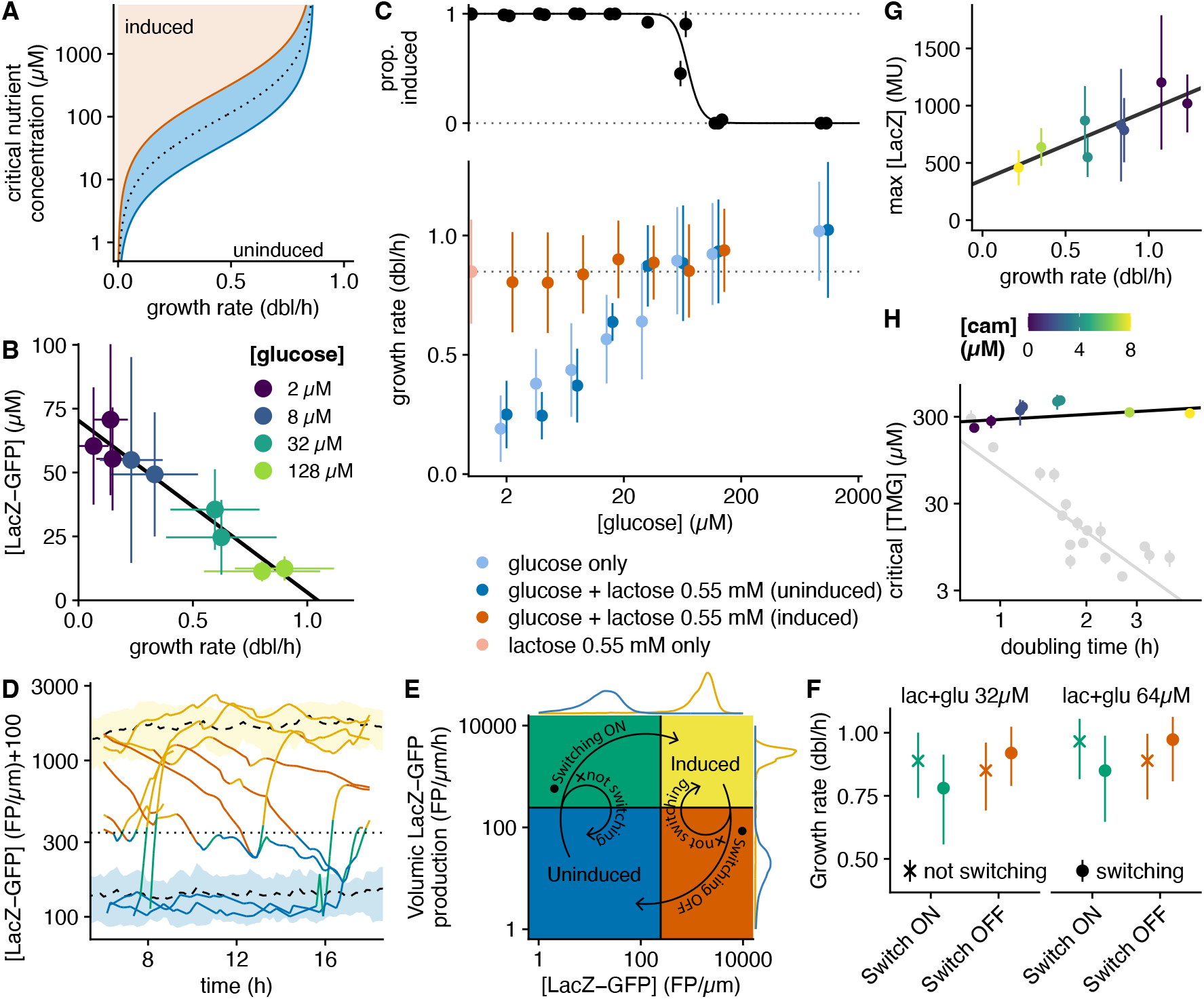
Growth-coupled sensitivity implements concentration-dependent sugar preferences **A**. Phase diagram of the *lac* operon regulatory switch when *lac* expression at full induction is regulated by CRP so as to decrease linearly with growth rate. The center of the bistable region (blue) perfectly tracks the inverted Monod function on lactose (dotted line). **B**. Mean and standard-deviations of LacZ-GFP concentration and growth rate in single cells grown in media with different concentrations of glucose (in colors) and induced with 200 µM IPTG. The linear fit shows that fully-induced *lac* expression decreases linearly with growth rate, as expected from CRP regulation. **C**. Fraction of single cells with induced *lac* operon when growing on mixtures of saturating lactose (0.55 mM) and different glucose concentrations (2 µM to 1.11 mM) (upper panel), and distributions of growth rates (mean ± s.d.) of the corresponding induced and uninduced subpopulations (lower panel, orange and dark blue; note that even when the fraction of induced cells is close to 1 or 0, enough cells remain in both states to measure the distribution of growth rates). For comparison, the distributions of single-cell growth rates on media with only glucose at different concentrations (light blue) and on media with only lactose (light orange) are also shown. Bacteria were grown in a modified version of the DIMM microfluidic device where media are flown through the growth channel during the whole experiment (1 to 3 independent replicates with more than 70 cells per condition; median ≈ 600 cells); no growth is observed in absence of carbon source (Fig. S12). **D**. LacZ-GFP concentration traces of individual lineages dynamically switching their *lac* operon on/off in a mixture of lactose (0.55 mM) and glucose (64 µM). Colors correspond to the *lac* expression state as described in E, and the horizontal dotted line shows the threshold between induced and uninduced cells (*e*^5.5^). Mean concentration ± standard-deviation of induced and uninduced cells is represented by the black dashed lines and yellow/blue ribbons respectively. For clarity, concentration values are offset by 100. **E**. On and off switches of the *lac* operon are defined as trajectories in the LacZ-GFP concentration (x axis) *vs* volumic production (y axis) parameter space. Cells switching on go from the blue quadrant (low concentration ; low production) to the yellow quadrant (high concentration ; high production), while cells switching off are defined inversely (arrows with a dot). Cells whose production goes above the threshold (resp. below) but whose concentration remains below the threshold (resp. above) are not considered switching (arrows with a cross). Thresholds are based on distributions measured for induced cells (marginal distribution in yellow, lactose 0.55 mM) and uninduced cells (marginal distribution in blue, lactose 0.55 mM + glucose 1.009 mM). **F**. Comparison of the growth rate distributions of cells switching on/off *vs* cells not switching in the two glucose+lactose mixtures where growth rates of induced and uninduced cells are similar (lactose 0.55 mM + glucose 32 or 64 µM). Cell lineages observed in the green quadrant (low concentration ; high production, see E) and orange quadrant (high concentration ; low production, see E) are each sampled once, and grouped according to the lineage fate (shown as symbol); points and error bars show median and interquartile range. **G**. Expression of the fully-induced *lac* operon as a function of growth rate when cells are exposed to subinhibitory levels of chloramphenicol at different concentrations (in colors). **H**. The critical level of TMG inducer is largely independent of growth rate when growth rate is modulated by chloramphenicol (colored points and black line for the linear fit). For comparison, critical TMG levels when growth rate is modulated by changing nutrients (same data as in Fig. 2F) is shown in light grey. In D and E, 2 to 8 µM of chloramphenicol were added to M9 + 0.2 % arabinose + 0.1 % casaminoacids; points and errors bars show the mean and standard error of three technical replicates; MU: Miller Units.

Remarkably, recent work in the context of general bacterial growth laws has shown that cAMP and CRP regulate the *lac* operon in precisely the manner of equation (1), i.e. the expression at full induction decreases linearly with growth rate, reaching zero at a growth rate λ_*_ [3].

These theoretical results suggest that *E. coli* may be exploiting GCS to switch between carbon sources in a concentration-dependent manner such that the resulting growth rate is maximized. For example, in contrast to the current view that *E. coli* always prefers to consume glucose when grown on a mixture of glucose and lactose, the theory predicts that whenever the glucose concentration is sufficiently low, i.e. when the growth rate that can be attained consuming glucose is lower than the growth rate that can be attained consuming lactose, the *lac* operon will be induced.

Unfortunately, it is not possible to test such predictions using batch culture experiments, because substantial reductions in growth rate only occur when the glucose concentration becomes so low that cell densities in batch cultures are too small to quantify using standard techniques. We thus turned to microfluidics for testing these predictions. To ensure a homogeneous growth environment and avoiding nutrient gradients that would otherwise arise within dead-end channels at low nutrient concentrations [28], we modified our DIMM microfluidic device such that growth media continuously flow through the channels in which the bacteria grow.

Using this device, we could grow bacteria at different glucose concentrations ranging from the usual 0.2 % (11.1 mM) down to 5500 times less, at a mere 2 µM. We observed a large range of growth rates (from 1.06 dbl/h down to 0.19 dbl/h) and first measured fully-induced *lac* operon expression at each glucose concentration by adding 200 µM IPTG (which, in contrast to TMG, readily enters the cell without LacY). We verified that average LacZ-GFP concentration decreases linearly with growth rate (Fig. 3B), confirming reports based on bulk measurements where the availability of glycolytic sugars was modulated by titrating their importers [3]. Moreover, although there is much uncertainty about the kinetic parameters of the *lac* system, it is noteworthy that the theoretically predicted values are consistent with our measured values of *y*_0_ ≈ 70*µM* and λ_*_ ≈ 0.72*h*^-1^ (Fig. 3B; see SI section 3). Second, we provide, to our knowledge, the first experimental confirmation that growth rate increases as a Monod curve with glucose concentration at the single-cell level (Fig. 3C, light blue symbols, and Fig. S11).

We then monitored growth and *lac* operon expression on mixtures of lactose and glucose, with glucose varying over the same range of concentrations and lactose at saturating concentration (0.55 mM i.e. 0.02 %) (Fig. 3C and Figs. S12 and S13). When glucose concentrations are very low, almost all cells induce their *lac* operon and grow at rates similar to growth on media containing only lactose. However, even though this is not visible in Fig. 3C, small fractions of cells with uninduced *lac* operon remain and we observed that their growth rates match the growth rates observed when growing on the same concentration of glucose only. As the concentration of glucose increases, the growth rates of the uninduced cells increases and, around a glucose concentration of 50 µM, the distributions of growth rates of the induced cells and uninduced cells become virtually identical. Remarkably, it is exactly at this critical concentration that we also see the fraction of induced cells drop sharply, and at higher glucose concentrations virtually all cells are uninduced (Fig. 3C). That is, these single cell experiments confirm our theory that, in contrast to the classical diauxie picture, there is no fixed sugar hierarchy but *E. coli* induces its *lac* operon in a concentration-dependent manner so as to always grow on the carbon source that maximizes growth rate.

In the conditions where growth rates on glucose and lactose are near equal, i.e. at 32 µM and 64 µM of glucose, our time-lapse microscopy measurements allow us to directly observe a significant number events in which cells stochastically switch from uninduced to induced or vice versa (Fig. 3D). Such stochastic switches go through two phases. When a cell switches away from the uninduced state (blue area in Fig. 3E) it goes through a phase where the production rate of LacZ-GFP is already high, but the LacZ-GFP concentration is still low (green area in Fig. 3E). If the high production rate persists, the cell will eventually move to the induced state with high LacZ-GFP concentration (yellow area in Fig. 3E) but sometimes cells will return to the uninduced state (i.e. move from the green area in Fig. 3E back to the blue area). Similarly, for induced cells to switch off, they pass through a phase where the production rate is already low but LacZ-GFP concentration is still high (orange area in Fig. 3E).

Our time-lapse measurements also allow us to track fluctuations in the instantaneous growth rates of cells. If, as our GCS theory assumes, induction of the *lac* regulatory switch depends on growth rate through dilution, then this predicts that stochastic switches from uninduced to induced are more likely to be preceded by a transient dip in growth rate, whereas switches from induced to uninduced are more likely to be preceded by a transient increase in growth rate.

To test this prediction, we identified all cells that moved into either the green or orange areas of Fig. 3E, stratified them by whether these cells eventually switched or returned to their previous state, and compared the growth rates of these two groups of cells. As shown in Fig. 3F, we see that the predictions of our GCS theory are borne out in the single cell data: cells that eventually switch on tend to have lower growth rates than those that return to the uninduced state, whereas cells that eventually switch off tend to have higher growth rates than those that return to the induced state (see also Fig. S14). We cannot think of another interpretation of these results than that transient growth rate fluctuations indeed contribute to stochastic switches of the *lac* regulatory switch. These results on the *lac* system suggest that *E. coli* could more generally exploit GCS to implement a strategy by which induction of operons for alternative carbon sources is both growth rate and concentration dependent, and tuned so that only the catabolic operon of the carbon source that is able to support the highest growth rate is induced.

Finally, although above we have considered changes in growth rate due to changes in nutrient quality, growth rate can of course also be modulated by other environmental factors, e.g. by stresses that inhibit replication or translation. While, as discussed above, optimal concentration-dependent carbon source preferences require that critical inducer concentrations increase with growth rate, this should ideally only apply when growth rate is modulated by nutrient quality. In contrast, if growth rate is modulated by other environmental factors, critical concentrations should ideally remain unchanged. Such different behaviors as a function of *how* growth rate is modulated might appear difficult to reconcile with GCS theory, because the dilution rate will be affected in the same way, independent of what environmental factors cause the change in growth rate.

One way to overcome this challenge is by modifying how the expression *y*_*h*_(λ) at full induction varies with growth rate depending on how growth rate is modulated. That is, while optimal concentration-dependent switching requires that *y*_*h*_(λ) decreases linearly according to equation (1) when growth rate is modulated by nutrient quality, we show in section 5 of the SI that if *y*_*h*_(λ) increases proportionally to λ when growth rate is modulated by other environmental factors, then the critical inducer concentration remains independent of growth rate. Remarkably, studies on resource allocation in the context of bacterial growth laws have established that CRP regulation exhibits exactly this behavior. In particular, when growth rate is modulated by translation inhibition using sublethal doses of chloramphenicol, the expression of CRP targets increases in proportion to growth rate as shown in [3] and [29].

To test these final predictions of the GCS theory, we first used batch cultures and Miller assay to measure expression *y*_*h*_(λ) of the fully-induced *lac* operon as we modulated growth rate using translation inhibition, i.e. starting from the highest growth rate condition (arabinose plus cas amino acids) and adding increasing levels of chloramphenicol. We confirm that *y*_*h*_(λ) increases approximately linearly with growth rate (Fig. 3G), although in contrast to [3] and [29], our measurements exhibit a nonzero offset at LacZ levels at zero growth rate. Since direct proportionality was reported not only for CRP targets in both [3] and [29], but also for constitutive genes in [2], we attribute this moderate quantitative discrepancy to imperfections of our Miller assay experiments.

We next used the same experimental approach as described above to measure *lac* promoter activity during balanced exponential growth, and confirmed the prediction that the critical external concentration of TMG is indeed independent of the growth rate in this case (Fig. 3H and Figs. S15 and S16). This confirms that CRP regulation not only optimally tunes the growth rate dependence of critical external inducer concentration under changes in nutrient quality, but also ensures that this critical concentration remains unchanged when growth rate is modulated by translation inhibition.

## Discussion

While it has been observed previously that growth rate can affect the functioning of synthetic gene circuits [30,31], we here systematically studied how growth rate can affect the responses of regulatory circuits to environmental signals, focusing on the behavior of gene regulatory switches. Using simple mathematical models we have shown that, by setting the rate of dilution of intracellular molecules, growth rate generally affects the sensitivity of gene regulatory circuits and that, by tuning the parameters of these circuits, natural selection can exploit this generic growth-coupled sensitivity (GCS) in a variety of ways. In general, GCS allows cells to respond to external signals in a context-dependent manner, e.g. to only respond to a particular signal when growth rate is below a critical value, or to scale the critical level of an external signal with growth rate (Fig. 1).

In the simplest default scenarios, gene regulatory switches decrease their sensitivity to their inducers with growth rate. Indeed, using as a example a mutant variant of the *lac* operon in *E. coli* that is insensitive to cAMP-CRP regulation, GCS theory predicts that critical inducer levels increase approximately quadratically with growth rate and we verified these predictions experimentally (Fig. 2). However, this behavior can be altered or fine-tuned by additional growth rate-dependent regulation. For example, it has been shown that while the maximal expression of carbon catabolism operons such as the *lac* operon decreases with growth rate when growth rate is modulated by nutrient quality, it increases proportional to growth rate when growth rate is modulated through translation inhibition [3, 29]. Our GCS theory predicts that this causes critical inducer levels to be independent of growth rate when growth rate is modulated by translation inhibition, and we experimentally confirmed this prediction as well (Fig. 3H).

In addition, in contrast to the currently predominant view that there is a fixed hierarchy of sugar preferences, our theoretical analysis suggests that natural selection can exploit GCS to implement optimal concentration-dependent sugar preferences. In this general strategy, each alternative carbon source has its own regulatory switch consisting of a positive feedback loop coupled to its corresponding inducer, and all feedback loops are coupled to growth rate through dilution. When multiple carbon sources are present at different concentrations, GCS ensures that only the positive feedback loop of the carbon source that (at its current concentration) provides the highest growth rate will be super-critical, and the positive feedback loops of all other carbon sources will be automatically switched off (Fig. S17). In this way, only the catabolic genes for the sugar that maximizes growth rate at these concentrations will be switched on.

We derived that this requires that cAMP-CRP signaling fine tunes the full-induction expression of catabolic operons of different carbon sources to linearly decrease with growth rate. Strikingly, for the *lac* operon precisely such a relationship has been observed previously [3], and we here confirmed that this relationship is consistent with the theoretically predicted optimal relationship (section 3 of the SI and Fig. 3B). In addition, using single-cell experiments with mixtures of glucose and lactose, we confirm that the critical concentration for *lac* operon induction is exactly where the single-cell distributions of growth rates on glucose and lactose match (Fig. 3C). Moreover, at glucose/lactose mixtures where the growth rates on glucose and lactose are very close, we find that stochastic ‘on’ and ‘off’ switches of the *lac* operon in single cells are accompanied by exactly the growth rate fluctuations predicted by our GCS theory.

It will be interesting to explore to what extent this regulatory strategy also applies to other sugar mixtures. In particular, it is well-known that certain sugar mixtures are not used sequentially but in parallel [24], and we hypothesize that such situations might correspond to parameter regimes in which multiple regulatory switches are super-critical in parallel.

Although it is tempting to speculate that evolution may have tuned CRP regulation to specifically implement the optimal concentration-dependent sugar preferences that we observed here, it should be noted that the theoretical models that have been proposed to explain the so-called bacterial growth laws argue that the growth rate dependence of CRP regulation and even the Monod equation itself follow from the necessity to balance catabolic and anabolic intracellular fluxes [2, 3]. It is thus conceivable that the optimal concentration-dependent regulation of carbon source preference reported here is in fact an emergent property of regulatory switches operating within the context of these growth laws. If this is indeed the case, this would constitute a remarkable example of adaptive behavior emerging from more basic physical constraints.

Apart from the specific regulation of carbon source preferences that we investigated here, the fact that GCS causes regulatory circuits to typically become less sensitive with increasing growth rate is likely adaptive in general for bacteria. That is, GCS causes fast growing cells to stabilize their current state by effectively muting their response to fluctuations in external signals and causes slowly growing cells to become highly sensitive to external signals. This view is consistent with recent work from our lab that shows that gene expression noise in *E. coli* results to a large extent from the propagation of noise through the gene regulatory network, and that noise levels systematically decrease with growth rate [32]. This suggests a general strategy in which GCS causes slowly growing cells to more actively explore alternative gene expression states and we show elsewhere that such behavior is highly adaptive for bet-hedging strategies [33].

While we have here focused on the behavior of gene regulatory switches in bacteria, the coupling of gene regulatory circuits to growth rate through dilution is so general that it likely affects the operation of gene regulatory circuits across organisms and GCS might also play a role in development and cell differentiation in multicellular eukaryotes. For example, since even a transient decrease in growth rate can cause regulatory switches to induce, it is conceivable that modulation of growth rate could be used in development to induce particular cell fate commitments. Indeed, it seems that a mechanism of this type acts to control lymphoid and myeloid differentiation in mouse [34] and behavior suggestive of this mechanism has been observed for the commitment to neurogenesis of neural progenitors [35, 36]. Exploring how GCS may have been exploited in the regulatory circuitry implementing the development and cell differentiation in multicellular eukaryotes is a fascinating area for future study.

The insights of our GCS theory may also have important applications in biotechnology, i.e. for the design of synthetic circuits. For example, the theory elucidates how different regulatory switches are globally coupled through dilution rate, and this insight might be exploited to design circuitry so as to induce the desired regulatory switches in a growth rate dependent manner.

## Acknowledgments

We thank Diana Blank for performing all experiments with the Miller assay. Strain ASC662 was kindly provided by Daan Kiviet and Sander Tans, and strain U486 by Anat Bren and Uri Alon.

## Notes

### Competing Interest Statement

The authors have declared no competing interest.

### Summary of Updates

Manuscript revised to address reviewers' comment.

