## Supplementary Information for "Growth rate controls the sensitivity of gene regulatory circuits"

#### 1 Growth-coupled sensitivity in regulatory switches

##### 1.1 A minimal growth-sensitive switch: the auto-activating operon

The simplest possible regulatory switch consists of nothing more than an 'auto-activating TF', i.e. an operon containing a TF that positively regulates its own transcription (Fig. 1A). This switch does not couple to any internal or external signal and the switching solely occurs because of the coupling to growth rate through dilution. Since we are predominantly interested in characterizing the overall behavior of such a system, and not aiming to model single-cell stochastic behaviors, we will model this system using simple ordinary differential equations.

Although the operon may contain multiple genes, we focus on modeling the concentration $X$ of the TF. Instead of modeling transcription and translation explicitly, we follow the standard modeling approach, e.g. [14, 15], and assume that the rate of production of the TF is a simple Hill function of the TF's concentration $X$. We then obtain the following differential equation for the TF's concentration $X$:

$$\frac{dX}{dt} = a\left[\frac{1 + r\left(\frac{X}{X_0}\right)^h}{1 + \left(\frac{X}{X_0}\right)^h}\right] - (\lambda + \mu)X. \tag{S1}$$

In this equation $a$ is the basal rate of production of the TF when its concentration is very low, and $ar$ is the rate of production at full induction, i.e. when the TF's concentration $X$ is very high. That is, the TF activates its own expression by $r$-fold. The concentration at which the TF is produced at half its maximal rate is $X_0$, and $h$ is the exponent of the Hill function. The concentration $X$ decays due to both decay at rate $\mu$ and dilution at the growth rate $\lambda$.

The possible steady-state levels of the concentration $X$ are solutions of the equation

$$\frac{a}{\lambda + \mu} = G(X, X_0, r, h) = X\left[\frac{1 + \left(\frac{X}{X_0}\right)^h}{1 + r\left(\frac{X}{X_0}\right)^h}\right], \tag{S2}$$

where the dependence on growth rate is on the left-hand side of the equation, and the right-hand side depends on the concentration $X$ and the parameters of the Hill function $X_0$, $r$, and $h$.

For values of the Hill coefficient $h \geq 2$ this system exhibits bistability. The expression within the square brackets on the right-hand side of equation (S2) approximately equals 1 for values of $X$ much smaller than $X_0$ and approximately $1/r$ for values of $X$ much larger than $X_0$. Thus, the function $G(X, X_0, r, h)$ is approximately $X$ for small $X$, reaches a maximum around $X_0$, then decreases, reaches a minimum, and finally grows as $X/r$ for large $X$.

Note that the left-hand side of equation (S2) is a decreasing function of the growth rate $\lambda$. Thus, at very high growth rates there is only one steady state given by the low value

$$X_l = \frac{a}{\lambda + \mu}, \tag{S3}$$

and, provided $a/\mu$ is sufficiently large, at very low growth rate there is a single steady state at the high value

$$X_h = \frac{ra}{\lambda + \mu}, \tag{S4}$$

with $X_h = rX_l$. In between there is a bistable regime whenever $a/(\lambda+\mu)$ is between the local minimum and local maximum of the function $G(X, X_0, r, h)$.

To keep the solution as simple as possible, we will assume $h = 2$, which can be interpreted as the TF binding as a dimer (although it may also be implemented in other ways, e.g. by a particular promoter architecture). For $h = 2$, the local extrema of $G(X, X_0, r, h)$ occur at $X$ values

$$X = X_0 \sqrt{\frac{r - 3 \pm \sqrt{r^2 - 10r + 9}}{2r}}. \tag{S5}$$

Note that real valued solutions only occur when $r > 9$, i.e. a fold-change of at least 9-fold. We define $G_{\min}(r)$ and $G_{\max}(r)$ to be the values of the function $G(X, X_0, 2, r)$ at its local minimum and maximum for a given value of $r$. Using this together with equation (S2), we find that the critical growth rates $\lambda_{\min}$ and $\lambda_{\max}$ that bound the bistable region are given by

$$\lambda_{\min} = \frac{a}{G_{\max}(r)} - \mu \tag{S6}$$

and

$$\lambda_{\max} = \frac{a}{G_{\min}(r)} - \mu. \tag{S7}$$

As an example of a biologically plausible growth-coupled auto-activating system we imagine a phage operon that activates its own expression at a critical concentration of $X_0 = 100$ TFs per cubic micrometer, a fold-change of $r = 500$, and a decay rate $\mu = 0.013$ per hour, i.e. the protein's half-life is about 53 hours. We note that the behavior of the model is independent of the units of protein concentration $X$ and time. For definiteness, we will choose hours for the unit of time and molecules per cubic micrometer for the units of TF concentration.

Fig. 1B shows the phase diagram obtained with these parameters as a function of the basal production rate $a$ and the cell's doubling time $t_d = \log(2)/\lambda$ with the white region corresponding to parameters for which the operon is stably off, the orange region corresponding to parameters where the operon is stably on, and the blue region corresponding to parameters where the operon is bistable.

For a range of intermediate basal expression levels $a$ the system goes from stably off, to bistable, to stably on as the growth rate is decreased. If we set $a = 0.061$ molecules per cubic micrometer per hour, the system is bistable for doubling times between 5 and 50 hours (dashed line in Fig. 1B).

Figure 1C shows the induction curve of the auto-activating operon as a function of doubling time, i.e. it shows the steady-state operon expression $X$ as a function of the doubling time, with the solid lines showing the stable steady states and the dashed line indicating unstable steady states. We see that when the cells are doubling faster than every 5 hours, the steady-state number of TFs per cubic micrometer (which is roughly the volume of a single cell) is less than 1 and the system is stably off. When the cells double less than once every 5 hours a second stable state appears with over 100 TFs per cubic micrometer. Once the cells double less than once every two days (50 hours) the low expression state disappears and all cells are guaranteed to switch to a high expressing state with on the order of a thousand TF molecules per cubic micrometer. It is conceivable that, using such an auto-activating operon, a phage could implement a strategy where it remains integrated in the genome as long as the host cell keeps growing and dividing, but switches to a lytic state once the growth rate falls below a critical value (here a doubling of once every two days). For slow growth, i.e. doubling between every 5 and 50 hours, the phage population would be expected to behave stochastically, with some lysing, and some remaining integrated.

#### 1.2 Signal dependent decay of the auto-activating protein

All that is needed to turn the minimal auto-activating system into a typical bistable switch with growth-coupled sensitivity is to couple the auto-activating TF to an external signal $s$, and this can be done in a number of different ways. For example, one can make the signal $s$ positively regulate the decay of the TF (Fig. 1D). Such a scheme is used in the core circuitry of the competence system in B. subtilis [12]. Here the ComK TF auto-activates itself while it is degraded by a specific ClpCP/MecA protease complex. A key external signal in this system is quorum sensing. The quorum sensing signal activates expression of ComS, which competes with ComK for binding to the protease ClpCP/MecA. In this way, the decay rate of ComK depends negatively on the amount of quorum sensing $s$.

Let $C$ denote the decay rate of the auto-activating TF (i.e. ComK) when all the protease is unbound and free to bind ComK. Next we will quantify the strength of the quorum sensing signal by the rate $s$ at which the competitor ComS binds to free protease and denote the rate of unbinding of bound protease by $s_0$. Assuming that the binding and unbinding dynamics is fast relative to other timescales in the system, the decay rate $C(s)$ at quorum signal level $s$ is then

$$C(s) = \frac{s_0}{s + s_0} C. \tag{S8}$$

The steady-state equation for the concentration $X$ of the ComK TF now becomes:

$$\frac{a}{\lambda + \mu + \frac{s_0}{s_0+s} C} = X \left[ \frac{1 + \left(\frac{X}{X_0}\right)^h}{1 + r\left(\frac{X}{X_0}\right)^h} \right]. \tag{S9}$$

We see that this equation is almost exactly the same as the steady-state equation (S2) for the minimal system. The only change is that the growth rate $\lambda$ is replaced by the sum $\lambda + \frac{s_0}{s_0+s} C$ of growth rate and decay due to the protease. Consequently, the system will be kept off whenever either the growth rate is high, or the signal $s$ is low. Only when both $s$ is large and the growth rate is low will the total decay rate be small enough for the positive feedback to set in. This is of course exactly the kind of behavior one would want the competence system to exhibit.

Like before, the bistable regime occurs when the left-hand side of equation (S9) lies between $G_{\min}(r)$ and $G_{\max}(r)$. Solving for the critical amount of signal

$s$ as a function of growth rate $\lambda$ and fold-change $r$ we find for the signal strengths that bound the bistable regime:

$$\frac{s_{\min}}{s_0} = \frac{C}{\frac{a}{G_{\min}(r)} - \lambda - \mu} - 1, \tag{S10}$$

and

$$\frac{s_{\max}}{s_0} = \frac{C}{\frac{a}{G_{\max}(r)} - \lambda - \mu} - 1, \tag{S11}$$

where $G_{\min}(r)$ and $G_{\max}(r)$ are again the values of the function on the right-hand side of equation (S9) at its local minimum and maximum and these depend only on the structural parameters of the promoter, i.e. the critical TF concentration $X_0$, the fold-change $r$, and the Hill coefficient $h$.

As an example, we took the same parameter settings as for the auto-activating TF of the previous section, i.e. $X_0 = 100$ proteins per cubic micrometer, $r = 500$, $h = 2$, $a = 0.061$ per cubic micrometer per hour, and $\mu = 0.013$ per hour. Furthermore, we set the half-life of the TF when all protease is free at 5 minutes, corresponding to $C = 8.32$ per hour. Fig. 1E shows the resulting phase diagram.

As in the simple auto-activating operon, this system can be induced by changing the growth rate of the system. Figure 1F shows the induction curves of the system as a function of doubling time at signal intensities $s = 10^4 s_0$ (in black) and $s = 100 s_0$ (in gray), corresponding to the black and gray dotted lines in the phase diagram of Fig. 1E. Note that, because at high signal intensity the protease is effectively inactive, the induction curve at high signal intensity is very similar to that of the auto-activating operon. At low doubling times the system is stably off and at a doubling time of 5 hours the system becomes bistable, with an 'on' state of more than 100 molecules per cubic micrometer. From a doubling time of 50 hours onward, the system is stably on. At the lower signal intensity of $s = 100 s_0$, the system remains in the off state longer and only becomes bistable at doubling times of about 12 hours or more. In addition, the system stays bistable no matter how large the doubling time becomes, i.e. even at growth arrest the system remains bistable.

Instead of keeping the signal strength $s$ fixed and varying growth rate, we can of course also keep the growth rate fixed and vary the signal strength. Note that, if the doubling time is less than 5 hours the system is locked in the off state independent of the strength of the external signal $s$. When the doubling time is between 5 and 50 hours, the system switches between locked in the off state and bistable when $s$ is over a critical value. This critical value is near $s = 60 s_0$ for large doubling times, and increases as growth rate increases. However, as long as the doubling time is less than 50 hours, the system remains bistable no matter how large the signal $s$, i.e. analogous to the behavior as a function of doubling time for $s = 100 s_0$. For doubling times longer than 50 hours, there is another critical value in signal $s$ beyond which the system is locked into the on state.

Note that, for purpose of illustration, we only considered one parameter regime of the auto-activating operon coupled to a signal through the decay rate of the TF, and that many other parameter regimes are possible that might lead to qualitatively different behaviors. For example, we assumed that both binding and unbinding of the signal molecule to the protease are fast, but one could also explore models where this binding dynamics is not always fast compared to the dilution rate $\lambda$. In addition, we assumed that the overall protease concentration $C$ does not itself depend on growth rate, but depending on how the expression of the protease is regulated, its concentration may itself be a decreasing or increasing function of the growth rate $\lambda$. Different parameter regimes might lead to different behaviors and phase diagrams which can be exploited by natural selection to implement different desired behaviors of the circuit. However, exploration of all the possible parameter regimes of this circuit is beyond the scope of this work.

#### 1.3 Two-component systems: activating the TF through phosphorylation

Probably the most common class of a systems in which an auto-activating TF is coupled to an external signal is the class of two-component systems in bacteria that auto-activate their own expression. In these systems the operon in question contains both a response regulator TF and a membrane-bound kinase. The response regulator TF only binds DNA in its phosphorylated form (often as a dimer) which is controlled by the membrane bound kinase. Whenever a specific external signal binds to the extracellular receptor domain of the kinase, the kinase auto-phosphorylates its intracellular kinase domain and then transfers this phosphate to the response regulator. Thus, the fraction of the auto-activating TFs that are active (phosphorylated) becomes a function of the strength $s$ of the external signal (Fig. 1G). Examples of such an architecture include the PhoP/Q system that controls virulence in *Salmonella enterica*, the prmA/B system in *S. enterica*, the PhoB/R system in *E. coli*, the CusR/S system in *E. coli*, the KdpD/E system in *E. coli*, the VanR/S system in *Streptomyces coelicolor*, and the ComE/D system of *Streptococcus pneumoniae* [37, 38]. Of course, when studied in detail these systems tend to be quite complex and involve coupling to multiple signals, but they all share a core architecture of an auto-activating TF whose activity is controlled by the level of an external signal transduced by a membrane-bound kinase.

To model this general type of system we extend the minimal auto-activating operon model as follows:

- Instead of just a regulator at concentration $X$, the operon now also includes a kinase whose concentration we will denote by $Q$. For simplicity, we will assume that the translation and decay $\mu$ of both these proteins is equal, such that the total concentration of both these proteins is guaranteed to be equal, i.e. $X = Q$ (at least for this simple deterministic model that ignores stochastic fluctuations).

- There is an external signal that can cause the membrane-bound kinase to switch to an activated state and we will denote the concentration of kinases in the activated state as $Q_s$.

- We quantify the strength of the external signal by the rate $s$ at which inactive kinase molecules are activated and let $s_0$ be the rate at which activated kinase molecules switch back to the inactive state.

- Inactive TFs can, upon interacting with an active kinase molecule, switch to the activated (phosphorylated) state. We will denote the concentration of TFs in the phosphorylated state by $X_p$.

- We will assume that the rate at which an unphosphorylated TF switches to the phosphorylated state equals $kQ_s$ and that phosphorylated TFs switch back to the unphosphorylated state at a rate $k_0$.

- Phosphorylated TFs form complexes that positively regulate the expression of the X/Q operon. We again assume a Hill function with coefficient $h$ and assume that, when bound by the TF, the promoter expresses $r$-fold more than when not bound.

The differential equations for the *total* concentrations $X$ and $Q$ of TFs and kinases are identical and given by

$$\frac{dX}{dt} = a \left[ \frac{1 + r\left(\frac{X_p}{X_0}\right)^h}{1 + \left(\frac{X_p}{X_0}\right)^h} \right] - (\lambda + \mu)X, \tag{S12}$$

$$\frac{dQ}{dt} = a \left[ \frac{1 + r\left(\frac{X_p}{X_0}\right)^h}{1 + \left(\frac{X_p}{X_0}\right)^h} \right] - (\lambda + \mu)Q, \tag{S13}$$

where $X_0$ is again the concentration $X_p$ of active TFs at which the promoter is bound half of the time, $a$ is the basal rate of protein production, $\lambda$ is the growth rate, and $\mu$ the protein decay rate. Note that we have set up the model such that, when $X$ and $Q$ start at equal concentrations, they are guaranteed to remain equal.

For the active forms $Q_s$ and $X_p$ we have the differential equations

$$\frac{dQ_s}{dt} = s(Q - Q_s) - (\lambda + \mu + s_0)Q_s, \tag{S14}$$

and

$$\frac{dX_p}{dt} = kQ_s(X - X_p) - (\lambda + \mu + k_0)X_p. \tag{S15}$$

The steady-state equations are straightforward to solve. For the concentrations of the active forms we find:

$$Q_s = \frac{s}{s + s_0 + \lambda + \mu} Q, \tag{S16}$$

and

$$X_p = \frac{kQ_s}{kQ_s + k_0 + \lambda + \mu} X. \tag{S17}$$

If we use the previous equation and the fact that $Q = X$, we can rewrite this as

$$X_p = \frac{k\frac{s}{s+s_0+\lambda+\mu} X^2}{k\frac{s}{s+s_0+\lambda+\mu} X + k_0 + \lambda + \mu}. \tag{S18}$$

If we define the dimensionless constant $C$:

$$C = \left(\frac{k_0 + \lambda + \mu}{kX_0}\right)\left(\frac{s + s_0 + \lambda + \mu}{s}\right), \tag{S19}$$

then we can write for the steady-state level of phosphorylated TF

$$X_p = \frac{X^2}{X + CX_0}. \tag{S20}$$

If we substitute this into the steady-state equation for $X$ and define $Z = X/X_0$ (the TF concentration relative to its critical level $X_0$) we find

$$\frac{a}{X_0(\lambda + \mu)} = F(Z, r, h, C) = Z \left[ \frac{1 + \left(\frac{Z^2}{Z+C}\right)^h}{1 + r\left(\frac{Z^2}{Z+C}\right)^h} \right]. \tag{S21}$$

This steady-state equation is in fact very similar to what we found for the minimal model (S2) above. The right-hand side approximately equals $Z$ when $Z$ is very small, and equals $Z/r$ when $Z$ is very large. Therefore, as before, the steady states at very low and high $X$ are $X_l = a/(\lambda + \mu)$ and $X_h = ra/(\lambda + \mu)$, respectively. However, the critical value of $X$ is not at $X_c = X_0$, or equivalently $Z_c = 1$, but is given by the solution of the equation:

$$\frac{Z_c^2}{Z_c + C} = 1. \tag{S22}$$

This simple quadratic equation has one positive solution given by

$$Z_c = \frac{1 + \sqrt{1 + 4C}}{2}, \tag{S23}$$

from which we already note that as $C$ goes from values much smaller than 1 to values much larger than 1, the critical value $Z_c$ goes from occurring at 1 to occurring at $\sqrt{C}$. However, note also that the situation is more complex because we want to investigate the steady-state solution both as a function of the external signal $s$ (which effects $C$) and of the growth rate $\lambda$, which simultaneously effects both the left-hand side of (S21) and $C$ as well.

Even with the simplifications already made, there are still many possible parameter regimes of this system that we could study. Here we focus on just illustrating one scenario that is relatively simple to calculate through analytically. First, we will assume that the de-activation of the kinase, as well as the dephosphorylation of the TF occur on much faster time scales than dilution and decay of the proteins, i.e. we assume $(\lambda + \mu) \ll k_0$ and $(\lambda + \mu) \ll s_0$. The constant $C$ then becomes independent of growth rate and we have

$$C = \frac{k_0}{kX_0} \frac{s + s_0}{s} \equiv r_p \frac{s + s_0}{s}, \tag{S24}$$

where we have defined $r_p$ to be the ratio between the rate of dephosphorylation and phosphorylation when both the TF and active kinase concentration $Q_s$ equal the critical concentration $X_0$.

**The location of the bistable region** Second, in order to simplify the calculation of the critical signal strengths $s$ at which the system switches, we will assume that the Hill coefficient $h$ is very large. Figure S1 shows the function $F(Z, r, h, C)$ of equation (S21) for a fold-change of $r = 100$, three different values of $C$ and for either a low ($h = 2$, dashed) or high ($h = 40$, solid) Hill coefficient. As can be seen from Fig. S1, for large $h$ the function $F(Z, r, h, C)$ switches from $Z$ to $Z/r$ over a very narrow range in $Z$ around the critical point $Z = Z_c$. Thus, for high Hill coefficients, the bistable region occurs whenever the ratio $Z_0 = a/(X_0(\lambda + \mu))$ lies between $Z_c/r$ and $Z_c$.

Combining equation (S23) for $Z_c$ as a function of $C$ with equation (S24) for $C$ as a function of $s$ we can solve for the signal strengths $s_{\min}$ and $s_{\max}$ that bound the bistable regime and find

$$s_{\min} = s_0 \frac{r_p}{rZ_0(rZ_0 - 1) - r_p} \tag{S25}$$

and

$$s_{\max} = s_0 \frac{r_p}{Z_0(Z_0 - 1) - r_p}, \tag{S26}$$

where $Z_0 = a/(X_0(\lambda + \mu))$ corresponds to the relative concentration $X_l/X_0$ when the system is in its off state. Note that, in order for $s_{\max}$ to be positive, we need to have $Z_0 > 1$ and $Z_0(Z_0 - 1) > r_p$. Similarly, for $s_{\min}$ to be positive we need to have $rZ_0 > 1$ and $rZ_0(rZ_0 - 1) > r_p$. That is, whether the system can become bistable and stably switched on as a function of $s$ depends on the parameters only through $Z_0$, $r_p$ and $r$.

Figure 1H shows the phase diagram of the auto-activating two-component system as a function of doubling time and signal strength $s/s_0$ for parameter settings $a = 20$ molecules per cubic micrometer per hour, $\mu = 0.01$ per hour, $X_0 = 100$ molecules per cubic micrometer, a fold-change $r = 100$ and a relative dephosphorylation to phosphorylation rate of $r_p = 5$. At these parameters, the system is already bistable as a function of the signal strength even at a very high growth rate that corresponds to the maximum growth rate that *E. coli* is capable of. It is interesting to note that the onset of the bistable regime as a function of signal strength $s$, i.e. the bottom of the blue region, strongly depends on the growth rate. As an illustration, Fig. 1I shows the induction curves of the system as a function of signal strength at a 2 hour doubling time (gray) and at a 20 hour doubling time (black), corresponding to the gray and black dotted lines in Fig. 1H. We see that although the growth rate differs only by a factor 10, the signal strength at the onset of the bistable region differs by a factor of roughly 100. That is, the critical signal strength decreases quadratically with doubling time at low doubling times, and eventually saturates when the doubling time becomes comparable to the half life of the molecules. Note also that at the 2 hour doubling time the system remains bistable even for very high signal strengths, whereas at 20 hour doubling times the system becomes stably switched on for signal strengths larger than $s \approx 0.1 s_0$.

Although this phase diagram is illustrative of the behavior that this system exhibits at some parameter settings, it should be noted that by varying parameters the system can show a number of qualitatively different behaviors. Although exploring the full range of this model's behavior is beyond the scope of this work, we note one or two points. First, because we have assumed that the basal expression rate $a$ is independent of growth rate, the low and high steady-state expression levels $X_l$ and $X_h$ both scale inversely with growth rate. Thus, as the growth rate decreases the system not only becomes much more sensitive to the signal, the low expression state $X_l$ also increases significantly such that, at least for some parameter settings, the $X_l$ at very low growth rates can become almost as high as $X_h$ at high growth rates. If we assume that the basal expression level $a$ grows roughly linearly with growth rate, then the steady-state levels $X_l$ and $X_h$ become independent of growth rate, and in this regime the critical value of the signal $s$ would also become independent of growth rate.

It might thus seem that either both the steady-state levels and the critical signal strength depend on growth rate, or both are independent of it. However, this is not necessarily the case if we drop the assumption that the rate of inactivation of the kinase $s_0$ and the rate of dephosphorylation of the TF $k_0$ are large relative to the rates of growth and decay. For example, one could imagine that once the TF $X$ is phosphorylated and forms a complex that activates the operon, this activated form $X_p$ is highly stable, i.e. dephosphorylates at a rate that is slower than the rate of growth and decay. In such a regime the system can exhibit both steady-state levels that are independent of the growth rate, and a sensitivity to the signal $s$ that does depend on growth rate.

These qualitatively different behaviors in different parameter regimes demonstrate the versatility of the GCS framework, i.e. by tuning parameters of the circuits, natural selection can realize different desired ways to couple the behavior of the regulatory circuit to growth rate.

#### 2 Growth-coupled sensitivity in a simple model of the *lac* operon

To illustrate growth-coupled sensitivity of the *lac* operon we use the simple mathematical model of the *lac* operon introduced in Ozbudak *et al.* [9], but now explicitly take into account the dependence on growth rate in the cell. There are two variables: $x$ is the internal concentration of TMG inducer and $y$ is the concentration of the proteins of the operon that are induced by it, i.e. lacZ and lacY, whose concentrations we assume to be proportional to each other (lacA is ignored in our model). The assumption is that TMG is imported by LacY and that the production of LacZ/LacY is inhibited by the LacI repressor. The variable $R$ denotes the concentration of active LacI repressor. The inducer inhibits the repressor by binding to it and this dynamics is considered so fast that, at any intracellular concentration $x$ of the inducer, one can assume one reaches the equilibrium active repressor level at a time scale that is short relative to the other time scales in the system:

$$R = \frac{R_T}{1 + (x/x_0)^h}, \tag{S27}$$

where $R_T$ is the total amount of repressor, $x_0$ is the concentration of TMG at which half of the repressors are active, and $h$ is a Hill coefficient. For the *lac* system, it has been determined that $h \approx 2$ [21]. In order to keep the model as simple as possible, we will assume that the total amount of repressor $R_T$ does not depend on the growth rate of the cell, i.e. that $R_T$ is roughly constant as the growth rate of the cell is varied. This assumption is consistent with available proteomics data on expression of LacI in different growth conditions [39]. Note that, if LacI levels were to vary systematically with growth rate in a known manner, then we could easily incorporate this into the theory without affecting its qualitative features. However, as we show below, the quantitative predictions of the theory are consistent with those derived under the assumption that LacI levels are approximately independent of growth rate.

The differential equations for the concentrations $x$ and $y$ are given by

$$\frac{dx}{dt} = by - (\lambda + \mu_x)x, \tag{S28}$$

and

$$\frac{dy}{dt} = \frac{a}{1 + R/R_0} - (\lambda + \mu_y)y, \tag{S29}$$

where $\lambda$ is the growth rate, $\mu_x$ and $\mu_y$ are the decay rates of TMG and LacY, and $R_0$ is the repressor level at which the promoter is expressed at half its maximal rate. The variable $a$ denotes the rate of production of proteins of the *lac* operon when the operon is fully induced. Note that, although it has been shown that at very high concentrations, TMG can also enter the cell independent of LacY [40], we neglect this relatively slow amount of LacY independent import. Similarly, while it was also shown in [40] that steady-state inducer concentrations differ in strains with and without LacA, this detail is left out of our simple model.

It is known that, through regulation by the transcription factor CRP, the maximal production rate $a(\lambda)$ depends on the growth rate and in section 3 below we will explore in detail how this affects the behavior of the system. However, when cAMP-CRP activity is held constant, it has been observed that the production $a$ of the lac promoter is roughly independent of growth rate [22]. For the validation of the GCS theory (Fig. 2E and F), we first use a ΔcyaA ΔcpdA mutant in which we set the cAMP-CRP activity to a constant level. Accordingly, we first analyze the simple case in which the maximal rate of production $a$ is *independent* of the growth rate $\lambda$.

The variable $b$ in equation (S28) denotes the inducer import rate, i.e. the amount of inducer that is imported per unit of concentration $y$ of LacY. In general, the rate $b$ will grow as a hyperbolic function of the external concentration $c$ of the inducer, i.e.

$$b = b_s \frac{c}{c + c_s}, \tag{S30}$$

where $b_s$ is the import rate at saturating external concentration of the inducer and $c_s$ is the concentration at which the import rate is at half its maximal rate. Note that, as long as the transporter is not near saturation, the import rate $b$ is simply proportional to the external concentration of the inducer $c$, i.e. $b \approx b_s c/c_s$.

We also note that equations (S28) and (S29) are identical to the equations used in Ozbudak et al. when one identifies the ratios $a/(\lambda + \mu_y)$ and $b/(\lambda + \mu_x)$ with the variables $\alpha$ and $\beta$ in their equations. That is, while in the equations of Ozbudak et al. the dependence of growth rate is 'hidden' within the definition of the parameters $\alpha$ and $\beta$, in our equations (S28) and (S29) we have brought out the dependence on growth rate explicitly.

#### 2.1 Steady states

At steady state, the variables $x$ and $y$ satisfy

$$y = x\frac{\lambda + \mu_x}{b}, \tag{S31}$$

$$y = \frac{a}{\lambda + \mu_y} \frac{\left(\frac{x}{x_0}\right)^h + 1}{\left(\frac{x}{x_0}\right)^h + \rho}, \tag{S32}$$

where we have defined the fold-change $\rho = 1 + \frac{R_T}{R_0}$, which gives the ratio of production rates of Lac proteins between the fully induced and fully repressed states of the operon. Substituting equation (S31) into (S32), the steady-state equation for the intracellular TMG concentration $x$ becomes

$$\frac{ab}{x_0(\lambda + \mu_x)(\lambda + \mu_y)} = F(x/x_0, h, \rho) = \left(\frac{x}{x_0}\right) \frac{\left(\frac{x}{x_0}\right)^h + \rho}{\left(\frac{x}{x_0}\right)^h + 1} \tag{S33}$$

Note that the function $F(x/x_0, h, \rho)$, i.e. the right-hand side of equation (S33), depends only on the dimensionless relative inducer concentration $x/x_0$ and structural parameters of the promoter and its repressor, i.e. the fold-change $\rho$, and the Hill coefficient $h$. In contrast, the left-hand side depends only on the dimensionless effective parameter

$$C = \frac{ab}{x_0(\lambda + \mu_x)(\lambda + \mu_y)} \tag{S34}$$

When growing on lactose, we have observed that there are on average $3000 - 12000$ molecules of LacZ-GFP per cell [11]. We also know that, while growing on glucose, the majority of cells have no *lac* expression at all, so that the average number of molecules per cell must be on the order of just a handful [11]. From this it follows that the ratio between the fully induced and fully repressed *lac* operon is on the order of $\rho \approx 1000$, which is consistent with previous estimates for the *lac* operon [9,21].

Figure S2 shows the function $F(x/x_0, h, \rho)$ for a fold-change of $\rho = 1000$ and for Hill coefficients $h = 2$ (blue), $h = 4$ (orange) and a large Hill coefficient $h = 50$ (green). We see that, for small values of $x/x_0$, the function $F(x/x_0, h, \rho)$ approximately equals $F(x/x_0, h, \rho) \approx \rho x/x_0$ and for large values of $x/x_0$ the function equals $F(x/x_0, h, \rho) \approx x/x_0$. Consequently, when the constant $C$ of equation (S34) is very low, the only solution is at low $x$:

$$x_l = x_0 \frac{C}{\rho} = \frac{ab}{\rho(\lambda + \mu_x)(\lambda + \mu_y)}, \tag{S35}$$

and when $C$ is very large, the only solution is at high $x$

$$x_h = x_0 C = \frac{ab}{(\lambda + \mu_x)(\lambda + \mu_y)}. \tag{S36}$$

The corresponding levels of the operon expression (and LacY concentration) are

$$y_l = \frac{a}{(\lambda + \mu_y)\rho}, \tag{S37}$$

$$y_h = \frac{a}{\lambda + \mu_y}. \tag{S38}$$

Note that the fold-change $y_h/y_l$ between the low and high levels is $\rho$, independent of growth rate.

##### 2.2 Phase diagram as a function of inducer level and growth rate

Bistability occurs whenever the line at the value $C$ of equation (S34) intersects the function $F(x/x_0, h, \rho)$ in three places, i.e. whenever there are 3 steady states. For example, for $h = 2$ and $\rho = 1000$, the two dashed lines in Fig. S2 correspond to the maximal and minimal values of $C$ for which bistability occurs. That is, the boundaries of the bistable regime are given by the extrema of the function $F(x/x_0, h, \rho)$

For $h = 2$, the extrema of $F(x/x_0, 2, \rho)$ occur at

$$\frac{x_{\text{min}}(\rho)}{x_0} = \sqrt{\frac{\rho - 3 - \sqrt{\rho^2 - 10\rho + 9}}{2}}, \tag{S39}$$

and

$$\frac{x_{\text{max}}(\rho)}{x_0} = \sqrt{\frac{\rho - 3 + \sqrt{\rho^2 - 10\rho + 9}}{2}}, \tag{S40}$$

which are only real valued when $\rho > 9$, i.e. bistability only occurs when $\rho > 9$. If bistability exists, the bistable regime runs over the range $C \in [C_{\text{min}}(\rho), C_{\text{max}}(\rho)]$, given by

$$C_{\text{min}}(\rho) = F\left(x_{\text{max}}(\rho)/x_0, 2, \rho\right), \tag{S41}$$

and

$$C_{\text{max}}(\rho) = F\left(x_{\text{min}}(\rho)/x_0, 2, \rho\right). \tag{S42}$$

Note that these values only depend on the fold-change $\rho$. For the *lac* operon's value $\rho = 1000$ we find $[C_{\text{min}}, C\text{max}] \approx [63.2, 500.5]$. For large $\rho$, a good analytical approximation is $[C_{\text{min}}, C_{\text{max}}] \approx [2\sqrt{\rho}, \rho/2]$.

Using the definition of $C$ (S34) we obtain for the critical import rates $b$ that bound the bistable region:

$$b_{\text{min}} = \frac{C_{\text{min}}(\rho)(\lambda + \mu_y)(\lambda + \mu_x)x_0}{a}, \tag{S43}$$

and

$$b_{\text{max}} = \frac{C_{\text{max}}(\rho)(\lambda + \mu_y)(\lambda + \mu_x)x_0}{a}. \tag{S44}$$

Note that these critical import rates determine critical external inducer concentrations through equation (S30).

Finally, we define the middle of the bistable region as the geometric average of its start and end, which ocurs at $C_{\text{crit}}(\rho) = \rho^{3/4}$, and has a critical import rate

$$b_{\text{crit}} = \frac{C_{\text{crit}}(\rho)(\lambda + \mu_y)(\lambda + \mu_x)x_0}{a}. \tag{S45}$$

Note that, precisely at this critical import rate $b_{\text{crit}}$, the system has an unstable fixed point at $y = y_h/\sqrt{\rho} = y_l\sqrt{\rho}$, i.e. also precisely at the geometric average of the fully repressed and fully induced expression levels.

##### 2.3 Rough parameter estimates

To provide an example of the phase diagram that these results imply we gathered rough parameter values for the key quantities. However, it should be noted that, for many of these parameters, mutually inconsistent values can be found in the literature, sometimes differing by more than an order of magnitude. The absolute values that result are thus only a rough guide.

To give an idea of the challenges with determining parameter values, consider the critical TMG concentration $x_0$ at which half of the LacI molecules are inactivated. In Boezi & Cowie 1961 [41] a mutant strain without LacY is used and the assumption is made that, for this strain, internal and external TMG concentrations will equilibrate. The authors then measure LacZ induction as a function of external TMG concentration and conclude that $x_0 = 3.5mM$. In contrast, Barkley *et al.* [42] use an *in vitro* method to estimate $x_0 = 9\mu M$, a value that is almost 400-fold lower! However, careful reading of Boezi & Cowie 1961 [41] shows that these authors erroneously assumed that $x_0$ corresponds to the inducer concentration where LacZ is at half-maximum expression. But that is incorrect. The half-maximum occurs at $\sqrt{\rho}x_0$ which is about 30-fold higher for $\rho \approx 1000$. The correct estimate of $x_0$ from these data would thus be $110\mu M$, which is still more than an order of magnitude larger than the estimate of Barkley *et al.* [42]. In a more recent paper, Marbach & Bettenbrock 2012 also show induction curves of LacZ in a strain without LacY and our fits to this data suggest $x_0 \approx 50\mu M$ [40]. Thus, the *in vivo* data suggest the range $50 - 100\mu M$ and we will use the middle of this range, i.e. $x_0 = 75\mu M$.

As far as we have been able to determine, little is known about the decay rates of both the TMG inducer and the Lac proteins other than that they must be highly stable. For illustration, we assumed that the decay rates of both the inducer and Lac proteins are $\mu_x = \mu_y = \log(2)/48 \approx 0.014$ per hour, i.e. half-lifes of 48 hours. Note, however, that these values only affect the phase diagram when the growth rate $\lambda$ becomes as small as the degradation rate.

For the expression of the fully induced lac operon in the strain with constant cAMP-CRP activity, we used an external cAMP concentration $(1mM)$ at which growth rate on lactose is optimized and expression is similar to that in the wildtype [20], which corresponds approximately to $a = 10\mu M h^{-1}$.

Finally, estimates of the pumping rate $b$ as a function of external inducer concentration are equally problematic as the estimates of $x_0$ because they involve measurements with different strains in different conditions than those used here and values reported in the literature can vary by up to 3 orders of magnitude. The general form of the pumping rate is given by equation (S30). Several works suggest $b_s$ in the range $1000 - 3000$ molecules per LacY per minute [43], which corresponds to $0.6 - 1.8 \times 10^5 h^{-1}$ per LacY. The corresponding half-maximum value $c_s$ is reported to be in the range $500 - 1000\mu M$. We will use the values in the middle of these ranges, i.e. $b_s = 1.2 \times 10^5 h^{-1}$ and $c_s = 750\mu M$.

Using these values the phase diagram as a function of doubling time and external TMG concentration in $\mu M$ can be calculated and the results are shown in Fig. 2B.

Note that whenever the growth rate $\lambda$ is considerably larger than the decay rate $\mu_x$ and $\mu_y$ the critical inducer concentration increases quadratically with growth rate $\lambda$. As mentioned above, we tested this prediction using a strain with fixed cAMP-CRP activity so that the maximal production rate $a$ should be approximately independent of growth rate [22]. We find that, in this strain, the critical inducer concentration indeed increases approximately as a power law of growth rate, with exponent $2.4 \pm 0.4$ (Fig. 2E-F), confirming that the sensitivity of the regulatory switch of the *lac* operon is strongly growth rate dependent, with an approximately 100-fold change in critical concentration between the fastest and slowest growth rate observed here. We note that the critical TMG concentrations predicted by our simple model are about $5-10$ fold lower than what we observe experimentally in Fig 2E. Considering the simplicity of our model and the large uncertainty regarding the parameters, we feel that it is remarkable that the absolute values of the predictions are within an order of magnitude of what we observe experimentally.

Finally, to obtain the steady-state *lac* expression $y$ as a function of the inducer concentration $c$ at a given growth rate $\lambda$, i.e. as shown in Fig. 2C, we simply solved the polynomial equation:

$$y = \frac{a}{(\lambda+\mu_y)} \left[ \frac{\left(\frac{by}{(\lambda+\mu_x)x_0}\right)^2 + 1}{\left(\frac{by}{(\lambda+\mu_x)x_0}\right)^2 + \rho} \right], \tag{S46}$$

with $b$ given by equation (S30).

#### 2.4 Why possible alternative interpretations are implausible

In our model, the quadratic increase of the critical TMG concentration results from the fact that, the steady-state transporter level $y$ is determined by a balance between production and dilution at rate $\lambda$, and that the internal TMG concentration is in turn given by a balance between import at rate $by$ and dilution at rate $\lambda$. That is, for both these molecules it is assumed that dilution dominates the rate at which the intracellular concentrations decay. Since LacY proteins are stable relative to the typical doubling time of the cells, this assumption is very likely accurate for the LacY proteins. However, for the relatively small molecule TMG we cannot exclude that there could be active transport out of the cell at a rate that is comparable or even larger than the dilution rate. However, if we assume that active export of TMG dominates over dilution, then in order to match our experimental observations (i.e. that critical TMG concentration increases at least quadratically with growth rate), we would have to assume that this export *increases linearly with growth rate*, i.e. to mimick the dependence of dilution rate on growth rate. We are not aware of any galactoside transporter that could export TMG and whose concentration or activity increases with growth rate.

We also note that it has been proposed that the role of the LacA acyltransferase is to, by acetylating competitor substrates of LacZ, promote these competing substrates to be exported out of the cell [44]. This raises the question as to whether the actions of LacA explain an increasing export rate of TMG as a function of growth rate. However, like the other members of the *lac* operon, due to the increased dilution, the expression of LacA *decreases* rather than increases with growth rate. Consequently, the potential effects of LacA move in the wrong direction with growth rate.

Furthermore, if molecules like TMG are indeed actively exported from the cell, then this would also call into question the *in vivo* measurements that were used to determine $x_0$ and $b_s$, since the interpretation of those experiments relied on the assumption that such active efflux does not exist.

Finally, as detailed in section 5 below, our simple model also correctly explains the constancy of the critical TMG concentration when growth rate is modulated through translation inhibition. This results from the fact that, when growth rate is modulated through nutrient quality, the lac expression at full induction decreases inversely proportional to growth rate, whereas it increases proportional to growth rate when growth rate is modulated through translation inhibition. If the intracellular TMG decay were dominated by export instead of decay, this would require that the expression of the corresponding transporter must always increase approximately linearly with growth rate in these experiments, independent of whether growth rate is modulated by nutrient quality or translation inhibition. Such behavior would be in contrast to the behavior of most other genes.

In summary, although it of course cannot be excluded that the rate of decay of intracellular TMG is determined by export, this requires assuming the existence of an unknown TMG transporter that scales its activity with growth rate in such a way as to precisely mimick the effects of dilution, independent of how growth rate is modulated. However, given that dilution necessarily occurs at a rate that equals growth rate, and this effect of dilution correctly predicts all our experimental observations, we feel that this is by far the simplest and most plausible interpretation.

### 3 Optimal carbon source switching

We have seen above that the critical concentration $c$ of inducer that is needed to induce regulatory switches increases with the current growth rate of the cells. Here we explore whether this growth-coupled sensitivity could be exploited to optimize the regulation of carbon source preference. We imagine a scenario in which cells are currently growing at a rate $\lambda_0$ on some carbon source and that, at some point, a new carbon source appears in the environment at concentration $c$. At what critical concentration should the cells decide to induce the regulatory switch that controls the expression of the catabolic genes for metabolizing this new carbon source? The optimal response would be to only induce this regulatory switch if the concentration $c$ is high enough that, once cells are growing on the new nutrient, the final growth rate is *larger* than the current growth rate $\lambda_0$. That is, if we denote by $\lambda(c)$ the growth rate on the new nutrient which is available at concentration $c$, the optimal strategy would be to only induce the regulatory switch when $\lambda(c) > \lambda_0$.

As has long been known, the growth rate that can be achieved when growing on a given carbon source is a hyperbolic function of the concentration $c$ of the carbon source, also known as a Monod function [25]. That is, if $c$ is the external concentration of the carbon source, the growth rate $\lambda(c)$ takes the form

$$\lambda(c) = \lambda_c \frac{c}{c + c_0}, \tag{S47}$$

where $\lambda_c$ is the maximal growth rate that can be achieved on the carbon source, and $c_0$ is the concentration at which the growth rate is half of the maximal growth rate $\lambda_c$. Inverting this relationship, we can determine the minimal concentration $c(\lambda)$ that is required so that the cells achieve a growth rate of at least $\lambda$

$$c(\lambda) = c_0 \frac{\lambda}{\lambda_c - \lambda}, \tag{S48}$$

which is an increasing function of $\lambda$ that diverges at $\lambda = \lambda_c$. Thus, in an optimal strategy the function $c_{\text{crit}}(\lambda_0)$, corresponding to the critical concentration at which the regulatory switch induces as a function of the current growth rate $\lambda_0$, should match exactly the inverse Monod function (S48), describing the concentration necessary to achieve growth rate $\lambda_0$.

In the previous sections we have seen that, for regulatory switches with positive feedback, the critical external concentration of the inducer that is needed to induce the switch naturally increases with growth rate due to the effects of dilution. In particular, equation (S45) shows that the critical import rate of the inducer grows approximately quadratically with growth rate $\lambda$ when the maximal production rate $a$ is constant. However, it is well known that the *lac* operon is also regulated by the TF CRP and potentially other growth-rate dependent physiological factors. As a consequence, when growth rate is modulated by nutrient quality, the maximal rate of production $a$ is a function $a(\lambda)$ of the growth rate $\lambda$ itself.

This raises the question as to whether it would be possible to choose the function $a(\lambda)$ in such a manner that the critical concentration $c$ of the inducer exactly matches the inverted Monod equation (S48). More precisely, we ask whether it is possible to choose the function $a(\lambda)$ such that the center of the bistable regime of the regulatory switch as a function of growth rate $\lambda$ is equal to the inverted Monod curve (S48).

We start from equation (S45) describing the critical import rate and make two modifications. First we replace the constant production rate $a$ with the (unknown) function $a(\lambda)$. Second, while the intracellular concentration $x$ of artificial inducers is given by the balance of import and dilution plus decay, i.e $x = by/(\lambda + \mu_x)$, the natural inducer allolactose needs to be produced through isomerization of lactose by LacZ. We here assume that a fixed fraction $\gamma$ of the imported lactose is converted to allolactose, so that we have for the allolactose concentration $x = \gamma by/(\lambda + \mu_x)$ and note that, in the next section, we will investigate in detail under what conditions on LacZ metabolism this assumption holds.

Using this and equation (S30) for the import rate as a function of the external lactose concentration, we find that the critical external lactose concentration $c$ for inducing the regulatory switch is a solution of

$$b_s \frac{c}{c + c_s} = \frac{C_{\text{crit}}(\rho)(\lambda + \mu_y)(\lambda + \mu_x)x_0}{a(\lambda)\gamma}. \tag{S49}$$

We now ask whether it is possible to choose the function $a(\lambda)$ such that the solution for the critical concentration $c$ from equation (S49) is identical to the equation (S48) for the critical concentration to achieve a growth rate of at least $\lambda$. We find that this is indeed possible, and is obtained when the function $a(\lambda)$ satisfies:

$$a(\lambda) = \frac{C_{\text{crit}}(\rho)c_s\lambda_c x_0}{b_s c_0 \gamma} (\lambda + \mu_y) \left(\frac{\lambda + \mu_x}{\lambda}\right) \left(1 - \frac{c_s - c_0}{c_s} \frac{\lambda}{\lambda_c}\right). \tag{S50}$$

Since the expression of the *lac* operon at full induction is given by $y_h(\lambda) = a(\lambda)/(\lambda + \mu_y)$, this is equivalent to demanding that the maximal expression of the *lac* operon as a function of growth rate should obey

$$y_h(\lambda) = \frac{C_{\text{crit}}(\rho)c_s\lambda_c x_0}{b_s c_0 \gamma} \left(1 - \frac{c_s - c_0}{c_s} \frac{\lambda}{\lambda_c}\right), \tag{S51}$$

where we have assumed that the growth rate $\lambda$ is sufficiently large compared to $\mu_x$ such that $(\lambda + \mu_x)/\lambda \approx 1$ and note that this assumption will also be investigated in detail in the next section.

Previous work has shown that CRP regulates expression from the *lac* operon in such a manner that, at full induction, its expression $y_h(\lambda)$ falls approximately linearly with growth rate [3], i.e.

$$y_h(\lambda) = y_0 \left(1 - \frac{\lambda}{\lambda_*}\right). \tag{S52}$$

Remarkably, this is the same functional form as expression (S51) that is needed for optimal carbon source switching. Furthermore, we have experimentally confirmed that this relationship also holds at the single-cell level (Fig. 3B) and fitting a linear relationship to our data we find $y_0 \approx 70\mu M$ and $\lambda_* \approx 0.72h^{-1}$.

Our theory thus predicts that, for optimal carbon source switching, the parameters should satisfy:

$$\lambda_* = \lambda_c \frac{c_s}{c_s - c_0}, \tag{S53}$$

and

$$y_0 = \frac{C_{\text{crit}}(\rho) c_s \lambda_c x_0}{b_s c_0 \gamma}. \tag{S54}$$

At these parameters equations (S51) and (S52) become identical. As illustrated in Fig. 3A, at those parameter settings the bistable region of the regulatory switch perfectly tracks the inverted Monod curve (dashed line) as a function of growth rate $\lambda$.

Although the parameters of the *lac* system are quite uncertain, the growth rate on saturating lactose is approximately $\lambda_c = 0.6h^{-1}$, the half-saturation of LacY is at $c_s = 500 - 1000\mu M$, and the lactose concentration $c_0$ at which the Monod curve reaches half of the maximal growth rate has been estimated to be $c_0 = 70 - 100\mu M$ [45, 46]. If we use the midpoints of these ranges, i.e $c_s = 750\mu M$ and $c_0 = 85\mu M$, we find $\lambda_* = 0.68h^{-1}$. Remarkably, this theoretically predicted value is within 5% of the experimentally estimated value of $\lambda_*$.

Similarly, using $C_{\text{crit}}(\rho) = \rho^{3/4} \approx 178$ for $\rho \approx 1000$, the previously mentioned values for $c_s$, $c_0$, $b_s = 1.2 \times 10^5 h^{-1}$ and $x_0 = 75\mu M$, then we find that $y_0$ equals $70\mu M$ if $\gamma \approx 0.01$. That is, if about 1% of the lactose transported into the cell is converted into allolactose, which seems not implausible. That is, not only does the functional form of the expression $y_h(\lambda)$ of the fully induced *lac* operon as a function of growth rate match the functional form needed to implement optimal carbon source switching, also the absolute parameters of the experimentally determined relationship are consistent with known parameters for the *lac* system.

In summary, we find that by combining the regulation of the *lac* operon by CRP and other growth rate dependent factors with growth-coupled sensitivity, it is possible for cells to implement an optimal carbon source switching strategy such that cells only induce the metabolic enzymes for a carbon source when its concentration $c$ is high enough such that switching to this carbon source is guaranteed to increase growth rate. Thus, instead of implementing a fixed preference hierarchy for different carbon sources, this strategy allows cells to choose carbon source in a concentration-dependent manner, such that growth rate is maximized.

#### 4 Modeling lactose metabolism

For artificial inducers the intra-cellular inducer concentration is a balance between import at rate $by$ and dilution at rate $\lambda$, i.e. $x = \gamma by/\lambda$ and in our derivation of optimal carbon source switching we assumed that the same functional relationship holds for natural inducers. In particular, we assumed that the intracellular allolactose concentration $A$ has the form $A = \gamma by/\lambda$, with $\gamma$ a constant. Here we develop a model of LacZ metabolism to investigate under what conditions this assumption applies.

The reactions catalyzed by LacZ are highly complex and different models of the *lac* system use different levels of coarse graining of this system [47–49]. In our model, the reactions we take into account include pumping of lactose into the cell by LacY, binding of lactose to free LacZ, hydrolysis of LacZ-bound lactose, isomerization of LacZ-bound lactose into allolactose (which includes its release from LacZ), binding of allolactose to free LacZ, hydrolysis of LacZ-bound allolactose, and dilution of all intracellular molecules at rate $\lambda$.

We denote the intracellular lactose concentration by $L$, the intracellular allolactose concentration by $A$, the total intracellular LacZ concentration by $Z$, the LacY concentration by $Y$, the concentration of LacZ-lactose complexes by $C_l$, and the concentration of LacZ-allolactose complexes by $C_a$. The dynamics of the lactose concentration is given by

$$\frac{dL}{dt} = bY - k_{bl}L(Z - C_l - C_a) - \lambda L, \tag{S55}$$

with $b$ the rate of lactose pumping per LacY and $k_{bl}$ the rate of binding of lactose to free LacZ.

The dynamics of the concentration of LacZ-lactose complexes is given by

$$\frac{dC_l}{dt} = k_{bl}L(Z - C_l - C_a) - (k_{hl} + k_i + \lambda)C_l, \tag{S56}$$

where $k_{hl}$ is the rate of lactose hydrolysis and $k_i$ is the rate of isomerization of lactose into allolactose.

The dynamics of the allolactose concentration is given by

$$\frac{dA}{dt} = k_iC_l - k_{ba}A(Z - C_l - C_a) - \lambda A, \tag{S57}$$

where $k_{ba}$ is the rate of allolactose binding to free LacZ.

Finally, the dynamics of the concentration of LacZ-allolactose complexes is given by

$$\frac{dC_a}{dt} = k_{ba}A(Z - C_l - C_a) - (k_{ha} + \lambda)C_a, \tag{S58}$$

where $k_{ha}$ is the rate of allolactose hydrolysis.

The steady-state complex concentrations are given by:

$$C_l = Z\frac{L/L_0}{1 + L/L_0 + A/A_0}, \tag{S59}$$

$$C_a = Z\frac{A/A_0}{1 + L/L_0 + A/A_0}, \tag{S60}$$

where we have defined the half-saturation concentrations

$$L_0 = \frac{k_{hl} + k_i + \lambda}{k_{bl}}, \tag{S61}$$

and

$$A_0 = \frac{k_{ha} + \lambda}{k_{ba}}. \tag{S62}$$

When we substitute the steady-state solutions for $C_l$ and $C_a$ into the equations for the dynamics of $L$ and $A$ we obtain the following steady-state equations:

$$bY - k_{bl} L \frac{Z}{1 + L/L_0 + A/A_0} - \lambda L = 0, \tag{S63}$$

and

$$k_i Z \frac{L/L_0}{1 + L/L_0 + A/A_0} - k_{ba} A \frac{Z}{1 + L/L_0 + A/A_0} - \lambda A = 0. \tag{S64}$$

To bring out the dimensionless structure of these equations, we rewrite them in terms of the dimensionless quantities

$$x = L/L_0 + A/A_0, \tag{S65}$$

$$f = \frac{L/L_0}{L/L_0 + A/A_0}, \tag{S66}$$

and the dimensionless parameters

$$Q = \frac{bY}{(k_{hl} + k_i + \lambda)Z}, \tag{S67}$$

$$d_l = \frac{\lambda}{k_{bl} Z}, \tag{S68}$$

$$p_i = \frac{k_i}{k_{ha} + \lambda}, \tag{S69}$$

and

$$d_a = \frac{\lambda}{k_{ba} Z}. \tag{S70}$$

Note that $x/(1 + x)$ is the fraction of LacZ that is in complex with either lactose or allolactose and $f$ is the fraction of the complexes that are bound by lactose. Further, the parameter $Q$ is the ratio of import of lactose and the maximal rate of lactose decay (through either hydrolysis, isomerization or dilution) when all LacZ molecules are bound by lactose. The parameter $d_l$ is the ratio between lactose dilution and the maximal rate of lactose binding to LacZ, when all LacZ is free. Similarly, the parameter $p_i$ is the ratio of production of allolactose through isomerization and the rate of decay of allolactose through either hydrolysis or dilution. And finally, $d_a$ is the ratio of the dilution rate of allolactose and the maximal rate of allolactose binding to LacZ, when all LacZ is free.

In terms of these dimensionless parameters and variables, the steady-state equations are

$$Q - f \frac{x}{1 + x} - d_l f x = 0, \tag{S71}$$

and

$$\frac{p_i f - (1-f)}{1+x} - d_a(1-f) = 0. \tag{S72}$$

Unfortunately, we generally do not know what these four dimensionless parameters are and the formal solutions of the above equations are complex roots of a third order polynomial that provide little insight. As far as we are aware, only *in vitro* measurements are available for the dynamics of purified LacZ exposed to very large concentrations of lactose [50]. Interpreting these is however highly challenging since the metabolic dynamics of the LacZ enzyme are considerably more complex than our model assumes. For example, apart from free, bound by lactose, or bound by allolactose, the enzyme can also be in a state bound by glucose only, by galactose only, and even jointly by galactose and another molecule of lactose or allolactose. Moreover, all these binding reactions are reversible and both hydrolysis and isomerization can run in reverse as well. The enzyme can even catalyze formation of higher order sugars such as trisaccharides (when either lactose or allolactose binds to the enzyme complexed with galactose). Thus, a full characterization of the LacZ kinetics would require at least a dozen parameters and there is simply not enough quantitative data available to meaningfully infer all these parameters.

Moreover, we note that for other kinetic parameters, such as the pumping rate of LacY, there were orders of magnitude discrepancies between the parameters inferred from *in vitro* and *in vivo* measurements. Consequently, it is impossible to put values on the effective kinetic constants $k_{bl}$, $k_i$, $k_{hl}$, $k_{ba}$, and $k_{ha}$ of our model, or even their order of magnitude. We thus focus on deriving under what conditions on these parameters the intracellular allolactose concentration as a function of pumping rate $b$, LacY concentration $Y$ and growth rate $\lambda$ is consistent with optimal carbon source switching.

We first consider a parameter regime that allows for a relatively simple analytical solution. In particular, the steady-state equations simplify considerably in the limit that most LacZ is not in complex, i.e. $x \ll 1$ which occurs when isomerization and hydrolysis are fast relative to the binding of lactose and allolactose to LacZ. Keeping only terms to highest order in $x$ we then find for the steady-state allolactose concentration

$$A = bY \left(\frac{k_i}{k_i + k_{hl} + \lambda}\right) \left(\frac{k_{bl} Z}{k_{bl} Z + \lambda}\right) \left(\frac{1}{k_{ba} Z + \lambda}\right). \tag{S73}$$

Note that, if isomerization of lactose ($k_i$), hydrolysis of lactose ($k_{hl}$) and binding of lactose to LacZ are all fast relative to dilution, but binding of allolactose to LacZ is slow relative to dilution, then we have

$$A = \frac{bY}{\lambda} \frac{k_i}{k_i + k_{hl}}, \tag{S74}$$

which matches the form $A = \gamma bY/\lambda$ that we assumed for optimal carbon source switching with $\gamma = k_i/(k_i + k_{hl})$. Moreover, in order to get $\gamma \approx 0.01$ we need that hydrolysis of lactose is about hundred-fold faster than isomerization.

#### 4.1 Numerical analysis

Besides this simple analytical solution, we also investigated whether there are other parameter regimes for which the internal allolactose concentration matches optimal carbon source switching. To do this, we note that optimal carbon source switching requires that the critical external lactose concentration $c_{\text{crit}}(\lambda)$ as a function of growth rate $\lambda$ equals the inverse of the Monod equation $c(\lambda)$. At the critical external lactose concentration, the lac system will be exactly in the middle of its bistable region, which is characterized by having an unstable fixed point that is exactly at the geometric average of the fully induced and fully repressed states. That is, at the critical inducer concentration $c_{\text{crit}}(\lambda)$, the system must have an unstable fixed point at LacY concentration

$$y_{\text{crit}}(\lambda) = \frac{y_h(\lambda)}{\sqrt{\rho}} = \frac{y_0}{\sqrt{\rho}}\left(1 - \frac{\lambda}{\lambda_*}\right). \tag{S75}$$

In order for this fixed point to exist, the internal inducer concentration $x$ has to satisfy

$$\frac{1}{\sqrt{\rho}} = \frac{1 + \left(\frac{x_{\text{crit}}}{x_0}\right)^2}{\rho + \left(\frac{x_{\text{crit}}}{x_0}\right)^2}, \tag{S76}$$

which is equivalent to

$$x_{\text{crit}} = \rho^{1/4} x_0. \tag{S77}$$

Thus, in order to implement optimal carbon source switching, whenever the external lactose concentration matches the inverse of the Monod equation $c(\lambda)$ of equation (S48) and the LacY concentration matches $y_{\text{crit}}(\lambda)$ of equation (S75), then the internal inducer concentration should match $x_{\text{crit}}$, and this should hold independent of growth rate.

Note that when the external concentration is the inverse Monod concentration $c(\lambda)$ the parameter $Q$ is given by

$$Q_{\text{crit}} = \frac{Y}{Z}\left(\frac{b_s}{k_{hl} + k_i}\right)\left(\frac{c(\lambda)}{c_s + c(\lambda)}\right) = b_r\left(\frac{c_0}{c_s \lambda_c}\right)\frac{\lambda}{1 - \frac{c_s - c_0}{c_0}\lambda}, \tag{S78}$$

where we have defined

$$b_r = \left(\frac{b_s}{k_i + k_{hl}}\right)\frac{Y}{Z}, \tag{S79}$$

and we used equation (S48).

Similarly, at this critical unstable fixed point $d_a$ and $d_l$ are given by

$$d_a = \frac{\lambda}{k_{ba} Z_{\text{crit}}} = \frac{\lambda\sqrt{\rho}}{k_{ba} y_0 (1 - \lambda/\lambda_*)} \tag{S80}$$

and

$$d_l = \frac{\lambda\sqrt{\rho}}{k_{ba} y_0 (1 - \lambda/\lambda_*)}, \tag{S81}$$

where we have assumed that the concentration $Z$ of LacZ is equal to that of LacY.

We then proceeded as follows. We performed a large sweep of the kinetic parameters

- $b_r \in [0.01, 100]$,
- $p_i \in [0.1, 1000]$,

- $k_{ba} \in [0.05, 500](\mu M h)^{-1}$,
- $k_{bl} \in [0.05, 500](\mu M h)^{-1}$,

and for each parameter setting varied the growth rate $\lambda$ from $\lambda 0.05h^{-1}$ to $\lambda = 0.6h^{-1}$, where the latter is the growth-rate on saturating lactose. For each value of $\lambda$ we then set $Q$ to $Q_{\text{crit}}$ of (S78), $d_a$ to (S80), $d_l$ to (S81), and calculated the normalized steady-state allolactose concentration $A/A_0 = (1-f)x$ by solving equations (S71) and (S72) numerically.

Finally, we calculated the mean $\mu$ and standard-deviation $\sigma$ of $\log(A/A_0)$ across the range of growth rates $\lambda$ and set $A_0 = x_{\text{crit}} e^{-\mu}$ so that on average the allolactose concentration $A$ matches the desired critical concentration $x_{\text{crit}}$ of (S77). Note that when $\sigma = 0$, the allolactose concentration at the critical point always matches $x_{\text{crit}}$ across the entire range of growth rates, ensuring optimal carbon source switching. Thus, the lower $\sigma$ the better a parameter setting supports optimal carbon source switching.

We found that there are many possible parameter settings that have low $\sigma$, ensuring close to perfect optimal carbon source switching (Fig. S10). In particular, the blue line in Fig. S10 shows the best parameter setting which has $\sigma = 0.035$ and occurs when the ratio $p_i$ of isomerization and hydrolysis of allolactose is large, the binding rate $k_{ba}$ of allolactose is low, and $A_0 \approx 29\mu M$. However, there are many other possible parameter settings that also have low $\sigma$. For example, the red, green and cyan curves in Fig. S10 show parameter settings with the largest $A_0$, the lowest $p_i$ and highest $k_{ba}$, respectively, that still have $\sigma < 0.1$.

In summary, besides the analytically determined regime (S74) that occurs when most LacZ is not in complex, our numerical analysis shows that there are many other possible parameter regimes for which the critical external lactose concentration almost perfect tracks the inverse Monod equation as a function of growth rate, ensuring optimal carbon source switching.

#### 5 Induction of the regulatory switch when varying growth rate through translation inhibition

All the results above presume that the growth rate $\lambda$ is varied through changes in carbon source. However, the growth rate $\lambda$ can also be altered through the introduction of stresses, for example by inhibiting translation through treatment with chloramphenicol. Previous work has shown that LacZ concentration grows roughly proportional to growth rate under this treatment [3] and we here confirm that LacZ levels at full induction increase approximately linearly with growth rate (Fig. 3D).

We thus find that $y_h(\lambda) \approx y_0 \lambda$ and, equivalently, that the maximal production rate $a(\lambda)$ is given by

$$a(\lambda) = a_0(\lambda + \mu_y)\lambda \tag{S82}$$

when varying $\lambda$ through translation inhibition. If we substitute this expression for $a(\lambda)$ into the equations for the critical import rate we find

$$b_{\min} = \frac{C_{\min}(\rho)(\lambda + \mu_x)}{a_0 \lambda}, \tag{S83}$$

and

$$b_{\max} = \frac{C_{\max}(\rho)(\lambda + \mu_x)}{a_0 \lambda}. \tag{S84}$$

Notably, these expressions are almost independent of growth rate as long as the growth rate $\lambda$ is large compared to the decay rate of the inducer $\mu_x$. That is, while the sensitivity of the *lac* regulatory switch is highly dependent on growth rate when growth rate is varied through changes in nutrients, when growth rate is varied through translation inhibition, the sensitivity of the regulatory switch remains roughly constant. As shown in Fig. 3E, these predictions of the model are confirmed by our experiments.

#### 6 Bacterial strains and media

All strains used in this studies are derivatives of *E. coli* K12 MG1655. The strain used for microfluidic experiments is ASC662 (MG1655 lacZ-GFPmut2) [19], further characterized in [11]. For induction experiments in constant environment measured with the Miller assay, we used U486 (MG1655 ΔcyaA ΔcpdA) supplemented with 1mM cAMP (growth rate modulated by different sugars) and MG1655 (growth rate modulated by subinhibitory levels of chloramphenicol). For induction experiments in constant environment measured with lacZ-GFP fluorescence, we used U486 lacZ-GFPmut2 and U486 Δcrr lacZ-GFPmut2 supplemented with 1mM cAMP, and MG1655 lacZ-GFPmut2 produced from the parent strain of U486 (CGSC #6300). The integration of LacZ-GFPmut2 in the chromosome for these strains is described below.

All experiments were done using M9 minimal media (Sigma-Aldrich) supplemented with 2 mM MgSO4, 0.1 mM CaCl2, and sugars as indicated (typically 0.2% for glucose, lactose or lactulose and 0.4% for glycerol). TMG and IPTG were diluted from frozen stocks in water (at 0.1 M and 1 M respectively), chloramphenicol (Cam) from frozen stock in ethanol (0.1M).

All experiments were carried out at 37°C. Note that the melAB operon is not expressed at this temperature [51] so that it cannot interfere with the regulation of the *lac* operon.

In experiments where the critical concentration of inducer as a function of growth rate was measured using GFP signal, we used strains constructed from the MG1655 #6300 of Coli Genetic Stock Center (CGSC). In order to integrate lacZ-GFPmut2 (taken from ASC662) into the chromosome of MG1655 (CGCS #6300) and of U486, we used the scarless in-frame insertion method developed by Cianfanelli *et. al* [52] To do so, we purified the pFOK plasmid from the diaminopimelic acid (DAP)-dependent strain JKe201, grown in LB supplemented with 100$\mu$M of DAP and 50 $\mu$g/mL of kanamycin. We amplified the lacZ-GFPmut2-lacY region with Q5 PCR (New England Biolabs) using the primers oth21 (atgtaGCGGCCGCcaggaaacgccaataacatacag) and oth22 (atgtaCTCGAGgtaataagcgttggcaatttaaccg); flanking regions are composed of a spacer and a NotI/XhoI sequence). We conducted a restriction-ligation between the PCR product and pFOK plasmid using NotI and XhoI restriction enzymes (New England Biolabs). We transformed the ligation product into a DH5α λpir strain with electroporation, amplified it and purified it. We subsequently transformed JKe201 with the obtained ligation product and followed the protocol from [52], using MG1655 (CGCS #6300) and U486 as recipient strains. Note that the conjugation was done by concentrating the recipient and donor strain together and incubating them for 6h at 37°C on the edge of an agar plate.

Finally, the strain U486 Δcrr lacZ-GFPmut2 was obtained by P1 transduction of the U486 lacZ-GFPmut2 strain obtained from the protocol above with a lysate of the Δcrr strain of the Keio collection (JW2410) [53]. The integrations and deletions were checked in all strains using PCR and Sanger sequencing.

### 7 Microscopy and image analysis

An inverted Nikon Ti-E microscope, equipped with a motorized xy-stage and enclosed in a temperature incubator (TheCube, Life Imaging Systems), was used to perform all microfluidic experiments. The sample was fixed on the stage using metal clamps and focus was maintained using hardware autofocus (Perfect Focus System, Nikon). Images were recorded using a CFI Plan Apochromat Lambda DM ×100 objective (NA 1.45, WD 0.13 mm) and a CMOS camera (Hamamatsu Orca-Flash 4.0). The setup was controlled using µManager [54] and timelapse movies were recorded with its Multi-Dimensional Acquisition engine (customized using runnables). Phase contrast images were acquired using 100 ms exposure (CoolLED pE-100, full power). Images of GFP fluorescence (ex 475/35 nm; em 525/50 nm; bs 495 nm) were acquired using different exposure and excitation intensity (Lumencor SpectraX, Cyan LED) for different types of experiments as described below (see 8.1 and 11.1).

Image analysis was performed using MoMA [18] as described in its documentation [55]. Raw image datasets were transferred to a centralised storage and preprocessed in batch. Growth channels (GCs) were picked randomly for downstream curation using slightly different sampling schemes for the two series of mother machine experiments: for experiments on induction during transient growth arrest, at least 30 GCs were picked randomly for each experiment (after discarding channels with structural defects or no cells growing at the first switch of condition). For experiments on induction in sugar mixtures, the use of mother machine channels with shallow reservoirs on their sides (see 11.1) led bacteria to eventually grow inside these reservoirs when nutrients concentration was low (typically less than 100 µM glucose), after what image analysis with MoMA was not possible anymore; since the delay until bacteria grow inside reservoirs is variable between experiments and between GCs, for each experiment and each condition at least 10 GCs were picked randomly among GCs where cells do no grow early inside reservoirs and curated manually in MoMA.

MoMA's default post-processing was used in order to refine the measurements of total fluorescence. Fluorescence arbitrary units were converted to number of GFP molecules using the procedure and conversion factors described previously [18].

### 8 Microfluidic experiments on the induction during transient growth arrest

#### 8.1 Experimental procedure

Experiments on the induction during transient growth arrest were done using the Dual Input Mother Machine (DIMM) [18] controlled using a pressure controller (OB1 mk3, Elvesys), following a procedure described in detail in a previous study [11]. Here, cells were exposed to glycerol 0.4% for 8h and subsequently to either glycerol 0.4% or lactulose 0.2% for 12h. In order to minimize phototoxicity, images were acquired only every 6 minutes (instead of every 3 minutes; with Lumencor SpectraX's Cyan LED at 17% with ND4), and fluorescence excitation was decreased 5 times (400 ms instead of 2000 ms) when bacteria were exposed to lactulose and the induction threshold was changed accordingly in the corresponding analysis.

##### 8.2 Analysis of induction under transient growth arrest

For this experiment, it is critical to pick a TMG concentration slightly below the lower threshold of *lac* operon bistability, so that no induction happens in glycerol but that there is enough inducer to support growth on lactulose. We hence characterized induction by TMG for our strain (ASC662) growing exponentially in M9 + 0.4% glycerol. Cultures grown overnight from single colonies in M9 + 0.4% glycerol and diluted 100× in the same media supplemented with variable concentrations of TMG (50 µM, 25 µM, 12.5 µM, 6.25 µM). After 8 hours of growth, cultures were concentrated by centrifugation, plated on 1% agarose slabs (M9 without sugar), and imaged readily using the same settings as for the mother machine experiments. For each culture, the concentration of LacZ-GFP was measured in 400 bacteria as the average fluorescence per pixel (corrected for the 100 grey level offset of the camera in the dark) in a 5-pixels wide circle close to the cell center. A simple graphical analysis revealed that 20 µM is the lower threshold of *lac* operon bistability in these conditions (Fig. S3A).

As predicted, no induction happened in Mother Machine experiments with 20 µM TMG in M9 + 0.4% glycerol. To the contrary, when switched to M9 + 0.2% lactulose supplemented with 20 µM TMG, cells stop growing transiently and then induce their *lac* operon (Fig. 2D; Fig. S3B), similar to the phenomenology under a switch to lactose [11]. However we note that the behavior on lactulose is more complex than on lactose. First, even in the absence of TMG, a fraction of the cells exhibited some growth and weak induction of the *lac* operon when growing on lactulose (Fig. S4A-B), possibly due to the very high activity of CRP and associated leaky *lac* operon expression in these conditions. In addition, because of the low concentration of extracellular TMG, even induced cells have relatively low steady-state levels of *lac* operon expression, i.e. 2 to 4 fold lower than on lactose (Fig. S4A), and this made Lac protein expression limiting for growth as indicated by the strong correlation between instantaneous growth rate and LacZ-GFP level (Fig. S4C). In order to verify that our analysis of the fraction of induced cells was done at steady state, we computed the distribution of induction lags using a procedure described previously [11] with a higher induction threshold (+1000 LacZ-GFP molecules instead of +200, in order to distinguish expression due to TMG from the weak expression observed in lactulose without TMG); this indicated that almost all cells which will ultimately switch do so within 5 h (Fig. S3B) and motivated our choice to estimate the fraction of induced cells between 7 and 8 h.

Overall, whereas no induction was observed in glycerol at all, the coupling to growth in the case of lactulose lead to significant induction of the *lac* operon in the large majority (84%) of cells after 7 h (Fig. 2D; Fig. S4A).

### 9 Measuring TMG sensitivity at steady-state growth using the Miller assay

The *lac* operon induction was measured in bulk for cultures in balanced exponential growth where the growth rate was modulated either by using different nutrients (0.2% ribose, 0.2% succinate, 0.2% rhamnose, 0.2% pyruvate, 0.2% mannose, 0.2% glycerol, 0.2% arabinose, 0.2% arabinose + 0.1% casamino acids; referred to hereafter as "sugars experiments"), or by adding variable subinhibitory levels of Cam (2 to 8 µM) to M9 + 0.2% arabinose + 0.1% casamino acids (referred to hereafter as "chloramphenicol experiments"). The activity of the *lac* promoter at variable TMG concentrations was obtained using the Miller assay [56], which measures β-galactosidase enzymatic activity and was performed according to a protocol and analysis by Kuhlmann *et al.* [21] with changes as follows.

Overnight cultures of bacterial strains were grown to saturation in 3 mL M9 + 0.2% arabinose for sugars experiments, or in 5 mL M9 + 0.2% arabinose + 0.1% casaminoacids for Cam experiments. Cultures were diluted to 150 µL into 96-well plates (Greiner) and grown for 16 h in a humidity-controlled incubator (Cytomat 2, Thermo Fisher; shaking at 600 rpm with 1 mm radius). For each condition, M9 was supplemented with 8 different concentrations of TMG in order to cover the induction range, and three combinations of dilution factor and delay before starting incubation were used so as to ensure having cells in mid-exponential phase after 16 h. All cultures were grown at 37°C. $OD_{600}$ was measured every 20 min in a Synergy H1 plate spectrophotometer (Biotek) using an Orbitor RS (Thermo Fisher) for plate moving and delidding; the cell-doubling rate ($\lambda$) was calculated for each sample as the slope of log2($OD_{600}$) vs. time plot via linear regression analysis (Fig. S7, Fig. S15). When $OD_{600}$ of all samples reached 0.06 and 0.2 (corresponding to $OD_{600}$ 0.2 to 0.5 for a spectrophotometer with 1 cm light path), a 5× dilution of the sample in M9 without sugar nor Cam (0.1 mL in 0.4 mL) was transferred to a 2-mL 96 well polypropylene block containing 0.5 mL Z-buffer [56], 20 µL 0.1% SDS, and 40 µL chloroform. All samples were thoroughly disrupted by repeated agitation with a multichannel pipettor. 200 µL of each sample was transferred to a flat-bottom transparent 96 well plate (Greiner). 40 µL of phosphate buffer containing 4 mg/mL o-nitrophenyl-β-D-galactopyranoside (ONPG) was added to each well and $OD_{420}$ measurements were performed during 10 h in a Synergy H1 plate spectrophotometer (Biotek), at 1-minute intervals for the first 30 minutes, and at increasing intervals (2 min during 90 minutes, 5 min during 8 hours) thereafter. Samples were maintained at 30°C throughout incubation, and the reader was pre-heated before loading the plate. All precultures and assays were performed in triplicate during each experiment, and one to four experiments were performed per condition (typically two).

Contrary to Kuhlmann, *et al.* [21], we did not observe an extended regime of linear dependence of $OD_{420}$ on time but rather that the slope of this relationship first increases and then decreases between the start of the measurements and the maximum $OD_{420}$ value (Fig. S5A). In order to establish how to measure the β-galactosidase activity from such time series, we compared the activities estimated using different time windows (initial slope over 15 min, 30 min, 1 h, 2 h, 3 h, or maximal slope) from culture samples containing variable concentrations of β-galactosidase. Practically, we mixed a culture induced with

500 µM TMG and an uninduced culture (so as to vary up to 10,000-fold the concentration of β-galactosidase per unit biomass), and diluted those samples in order to obtain 3 different cell concentrations ($OD_{600}$ 0.8, 0.2 or 0.05). Since the apparent β-galactosidase activity is expected to increase linearly with the relative β-galactosidase concentration, we concluded that the activity is best measured using the initial slope ($s$) over 30 min, for concentrations as low as a few molecules of β-galactosidase (Fig. S5B). Note that, at low LacZ concentrations, the $OD_{420}$ sometimes started by decreasing during up to 1 h (Fig. S5A right panel), in which case this initial decrease was discarded from the analysis. β-galactosidase activity ($A$) was expressed in Miller Units (MU) according to the formula $A = (1000\ s)/(0.5\ OD_{600})$ (where $OD_{600}$ was adjusted to be equivalent to using a 1 cm light path) [56], and the *lac* promoter activity ($\alpha$) was estimated as $\alpha = A\ \lambda$, where $\lambda$ is the cell-doubling rate defined above (in unit of 1/hr) and measured in each well. This measure of the promoter activity is motivated by the fact that the enzyme β-galactosidase is very stable so that in the balanced exponential growth, its "turnover" is governed by dilution due to cell growth [21].

For each condition, the TMG induction curve was fitted to the function

$$\alpha = b\,\frac{1 + f(c/k)^m}{1 + (c/k)^m} \tag{S85}$$

where $c$ is the TMG concentration, $b$ the basal *lac* promoter activity, $f$ the induction fold-change, $k$ the critical inducer concentration, and $m$ the Hill coefficient (Fig. 2E, Fig. S6, Fig. S16). While all other variables are almost independent of the growth rate, the critical inducer concentration $k$ shows a clear positive correlation to growth rate when the growth rate is modulated by nutrients.

To fit the exponent for the dependence of the critical inducer concentration $k$ as a function of doubling time $t = 1/\lambda$, we used all the estimated doubling times and critical concentration pairs $(t_i, k_i)$ for all individual replicate measurements $i$ and fitted a linear relationship $y_i = ax_i + b$ to the logarithms ($x_i = \log[t_i], y_i = \log[k_i]$). Note that, from the Hill function fits, we obtained an error-bar $\sigma_i$ for the relative error of the estimated critical concentration $k_i$ of replicate $i$. We assume that the estimated $y_i$ differ from those predicted by the linear relationship by both Gaussian measurement noise of standard-deviation $\sigma_i$ and additional intrinsic deviation between model and experiment of standard-deviation $\sigma$. Given this model, the probability of the data given $a$, $b$, and $\sigma$ becomes

$$P(D|a,b,\sigma) = \left[\prod_i \frac{1}{\sqrt{2\pi(\sigma^2+\sigma_i^2)}}\right] \exp\left[-\sum_i \frac{(y_i - ax_i - b)^2}{2(\sigma^2+\sigma_i^2)}\right]. \tag{S86}$$

We use a uniform prior over both $a$ and $b$, a scale prior $d\sigma/\sigma$ for $\sigma$, marginalize analytically over $b$ and maximize the resulting posterior over $\sigma$. From the resulting posterior of $a$ (i.e. at optimal $\sigma$) we then finally obtain the best slope $a = -2.4$ with an error-bar (i.e. standard-deviation of the posterior distribution over $a$) of 0.4 (Fig. 2F).

In contrast to the strong dependence of the critical concentration on growth rate when growth rate is modulated by nutrients, no dependence on the growth rate was observed when it was modulated with subinhibitory levels of chloramphenicol (Fig. 3E).

### 10 Measuring TMG sensitivity at steady-state growth using LacZ-GFP fluorescence

The *lac* operon induction was also measured in bulk using fluorimetry with a LacZ-GFP at the native locus for cultures in balanced exponential growth where the growth rate was modulated by using different nutrients, i.e. M9 with 0.2% ribose, 0.2% pyruvate, 0.2% mannose, 0.2% glycerol, 0.2% arabinose, and 0.2% arabinose + 0.1% casamino acids. We took advantage of the fact that, in our hands, this assay is comparatively easier than the Miller assay, to measure induction in several different strains. In particular, we wanted to investigate the effects of native cAMP-CRP regulation and inducer exclusion on TMG sensitivity. We thus measured *lac* operon induction with TMG in U486 lacZ-GFP supplemented with 1 mM cAMP (i.e. the same strain as used for the Miller assay, to confirm consistency), in the wild type MG1655 lacZ-GFP strain (to assess the effect of CRP regulation), and in U486 Δcrr lacZ-GFP. Notably, the crr deletion removes inducer exclusion, although it is also reported to affect CRP signaling [57] [23].

Overnight cultures of bacterial strains were grown to saturation in 2 mL M9 + 0.2% arabinose. Cultures were diluted (2000 to 10000 ×) to 160 µL into black 96-well plates with clear bottom (Greiner). Plates were incubated at 37ºC with continuous double orbital shaking (548 cpm frequency, 'fast' speed) in a Synergy H1 plate spectrophotometer (Biotek), where $OD_{600}$ and fluorescence (Ex 470 nm, Em 525 nm) were measured every 4 min. For each condition, M9 was supplemented (in duplicate) with 8 different concentrations of TMG in order to cover the induction range and several independent biological replicates were obtained for each condition.

We first background-corrected each growth curve by estimating background OD and fluorescence as the average value in a 20 time point window centered on the observation with minimal OD/fluorescence and subtracted these background values from all observations. Next we identified a 'growth-segment' of each curve by finding the time point with maximum (background-corrected) OD, and going backward from this time point find the segments of time points with background-corrected OD less then $OD_{max}/3$, background-corrected OD larger than 0.01, and (background-corrected) fluorescence larger than zero. For quality control we performed simple linear regressions of log-OD and log-fluorescence against time for the resulting growth-segment and extracted their Pearson correlation coefficients. Among the 1600 growth curves, there was a small number (21) with anomalously little growth or poor Pearson correlations and we removed these by hand. Growth-segments with less than 21 time points were also removed. This left 1579 of the 1600 growth curves for further analysis.

Next, we extracted for the growth-segment of each growth curve an average growth rate, an error-bar on growth rate, an average log-ratio $y = \log(\text{fluo}/\text{OD})$ of background-corrected fluorescence and OD, and an error-bar on this log-ratio $y$ as follows. We move a sliding window of length 21 across the growth segment and for each window perform a linear fit of log-OD against time and log-fluo against time. From these we extract an estimated log-ratio fluo/OD $y$ with error-bar and a growth-rate $\lambda$ with error bar. We then combine the estimated values of $y$ and $\lambda$ from each window into one final average $y$ and $\lambda$ (with error-bars) for the growth segment.

Finally, for each combination of a strain, media and replicate, we extracted the estimated $y$ values and their error-bars for the series of growth segments with different TMG concentrations and then fit an induction curve exactly as described in section 9 using equations (S85) and (S86). The resulting fits are shown in Fig. S8. For each induction curve we extracted the estimated critical TMG concentration and the average and standard deviation of doubling time across the corresponding series of growth segments.

Figure S9 shows the observed relationship between doubling time and critical TMG level for the U486 strain with constant cAMP-CRP (top left), the wildtype MG1655 strain (top right) and the U486 $\Delta$crr strain without inducer exclusion (bottom left). Symbols of the same color denote different replicates for the same growth media. It is notable that there is quite some variability across replicates and that, as indicated by the large error-bars on the doubling times, growth rates are quite variable even within one replicate series. Nonetheless the results for both the U486 strain and the wildtype MG1655 strain are consistent generally consistent with our results from the Miller assay experiments and are consistent with a quadratic decrease of critical TMG levels as a function of doubling time (dotted black lines).

We found that deletion of crr strongly affected the growth of the strains and their reproducicibility. Observed growth rates where often quite different from the growth rates of the other strains in the same media, e.g. with *slower* growth on arabinose with cas amino acids than on arabinose. There was also very substantial variation in behavior across both technical and biological replicates, e.g. the large variation in growth rates in pyruvate. Although the critical TMG concentration still generally decreased with doubling time for the crr deletion, there is so much variation that no clear quantitative relationship can be inferred from this data.

Therefore, to quantify most directly the effect of inducer exclusion on the critical TMG concentration, we plotted the fold-change in critical TMG concentration caused by the crr deletion against the fold change in growth rate caused by this deletion (Fig S9), bottom right panel). We find that in some media (i.e. arabinose, ribose, and glycerol), there is little systematic effect on either induction threshold or growth rate. For mannose, the crr deletion leads to an increase in growth rate which is accompanied by an increase in critical TMG concentration, as predicted by our GCS theory. For pyruvate and arabinose with case amino acids, there is substantial variation in growth rate across replicates and the change in critical TMG concentration correlates well with the change in growth rate, again supporting that dilution rate is indeed a key determinant of the induction threshold. In summary, the results in Fig S9 show that inducer exclusion is not a key determinant of the induction threshold and instead support that growth rate is a indeed a key determinant of the induction threshold.

### 11 Microfluidic experiments on the induction in sugar mixtures

#### 11.1 Experimental procedure

**Microfluidic device fabrication** The microfluidic device used in this series of experiments is similar to the DIMM device described earlier [18], with the main improvement that GCs are not closed on one end but rather designed to act like filters, letting media flow through but retaining cells. Given the small diameter of *E. coli* (typically between 0.6 µm and 1 µm in conditions considered here), it is very challenging to manufacture using UV soft lithography a constriction in the channel which is small enough to prevent cells from passing through and/or to block the flow through the GC. We solved this problem by modifying the initial design in two complimentary ways: each individual GC is surrounded by a large and shallow reservoir (width: 3 µm, depth: 0.25 µm, hence requiring a third microfabrication layer) as introduded by Norman *et al.* [58], and the back end of every reservoir is connected, via two small channels of the same depth as the GCs, to a back flow channel. This back flow channel is itself connected to an outlet to which a negative pressure is applied, thus generating a flow through the GCs running from the main flow channel to the back flow channel. Altogether, this allows media to flow through the GCs and their flanking reservoirs during the whole experiment, creating a homogeneous environment across the GCs, as indicated by the fact that the cell growth rate doesn't depend on the cell rank in the GC even at our lowest glucose concentrations of a few µM (Fig. S12B). Notably, this is in contrast to experiments in classical mother machines where gradients in growth rate along the GC are observed, even at high concentrations of glucose (e.g. at 0.2%). Importantly, this filter-like design of the GCs also greatly facilitates the cell loading step, which doesn't require concentrating the exponentially growing culture anymore, as the flow from the main channel through the GCs naturally drives cells into the GCs. Finally, our new device consists of 8 independent series, allowing us to conduct up to 8 different media switches with up to 8 different strains in parallel. Due to space constraints, the mixing serpentines that existed in the original DIMM design had to be removed, so that only experiments with "hard" switches from one input media to another can be performed, i.e. precluding mixtures of the two input media.

Microfluidic masters were produced through soft lithography by micro-resist Gmbh; most of the data were produced using a single master with GCs and reservoirs of appropriate dimensions (GC 0.8-1.0 µm width × 0.7 µm height ; reservoirs 3 µm width × 0.25 µm height). Device preparation was performed as described in the first series of microfluidic experiments, with two notable changes: surface activation in the plasma cleaner was reduced to 10-30 sec (typically 10 sec) at 1300-1500 µm of Hg in order to prevent collapse of the reservoirs; the device was primed with BSA (10 mg/mL) from the back channel outlet and water from the overflow outlet. In these experiments, only one of the inlets at each "dial-a-wave" junction was punched, as we were not intending to perform condition switches.

**Culture conditions and flow control** Bacteria were streaked onto LB agar plates from frozen glycerol stocks stored at −80°C. Overnight preculture was grown from single colonies in M9 minimal medium supplemented with 0.2% of glucose. The next day, cells were diluted 100-fold into fresh medium and harvested after 4–6 h, at $OD_{600}$ 0.05-0.15.

In the microfluidic devices, bacteria were grown in M9 minimal media with three different types of treatment: either glucose only at variable concentration, or a mixture of lactose (0.2%) and glucose (at different concentrations), or glucose (at different concentrations) supplemented with 200 µM IPTG in order to inhibit LacI repression.

For this, the experimental setup was prewarmed and initialized. The primed microfluidic chip was mounted, connected to the media supply and flushed with running media for 30 min or more to rinse the BSA. Flow control was achieved using a pressure controller, applying 1500 mbar on the 8 media inlets, and -600 mbar on the back channel outlet. The resulting flow is approximately 4-5 µL/min through each flow channel. The exponentially growing bacteria culture was sucked ($\approx$ 300 µL) using a syringe connected to the cell outlet, and all 8 cell-containing tubing pieces were connected to the pressure controller through a manifold, and pressurized at 1800 mbar. This resulted in the cells being pushed into the flow channels of each series, and under the effect of the flow generated by the vacuum, the cells were aspirated in the GC very efficiently (15-30 min to load all 8 series). It was necessary to disconnect the cell-containing tubing from the manifold once the series has been properly loaded to avoid clogging the flow channels.

After loading, the acquisition was quickly started. We typically acquired 4 positions per series, comprising ca. 40 GCs each. Phase contrast was acquired every 3 min with 100 ms exposure time, while GFP was acquired only every 9 min with low intensity (100 ms exposure time with SpectraX Cyan LED at 48%) in order to minimize phototoxicity.

#### 11.2 Analysis of growth and induction

**Population level growth and induction** In experiments with a mixture of lactose (0.55 mM i.e. 0.02%) and glucose (at different concentrations), bacteria were pre-cultured in M9 + 0.2% glucose where they grow fast ($\sim$ 1 doubling/h) and the *lac* operon is repressed. Upon transfer into the mixture of glucose and lactose in the microfluidic device, they decrease their growth rate depending on the sugar concentrations and possibly induce their *lac* operon; note that in most conditions this induction is typically irreversible i.e. the cell and its progeny will remain induced. To analyse how the fraction of induced cells and the growth rate depend on the concentration of glucose in the mixture, we needed (1) to set a criterium for calling a cell induced and (2) to establish how long it takes for bacteria to adapt to the new condition so that we can measure variables at steady state. In the following, the single-cell growth rate is measured as the slope of the linear regression of log(length) vs. time.

We first established that bacteria cannot grow on nutrient traces coming from the media or the setup (i.e. the microfluidic device, tubing, etc.), as indicated by the distribution of growth rate of cells exposed to M9 minimal media without carbon source (measured over windows of 1 h; Fig. S12A). In order to pick a criterium for induction, we examined time series of fluorescence in single cells, as well as distribution of LacZ-GFP levels at different glucose concentrations (Fig. S12C) and observed that a threshold of 250 LacZ-GFP molecules clearly separates the two subpopulations at all glucose concentrations. In addition, we further checked that, using this threshold, virtually all cells were either induced or uninduced over their entire trace (0.8% of cells were induced more than 5% but less than 95% of their trace and these were discarded from further analysis).

Looking at the fraction of induced cells as a function of time, we found that it stabilised within at most 5 h, and hence discarded the first 5 h of each experiment. For concentrations of glucose close to the threshold of induction (32 and 64 µM), we noticed that it took longer to reach a stable fraction of induced cells and discarded the first 14 h instead. Doing so, we discarded two replicates which terminated before 14 h (Table S2, 32 µM glucose on 20210122 and 20210305). Moreover, examining LacZ-GFP expression level revealed that one replicate was an outlier (probably due to an experimental mistake) and it was hence discarded from further analysis (Table S2, 16 µM glucose on 20210122).

In order to focus on cells for which growth rate can be reliably estimated, we filtered out cells which were monitored for less than 10 consecutive time points (30 min); note that this limit is below the shortest cell cycle that can be observed in our conditions, even at saturating glucose concentration). The same filtering criteria were applied to measure the distributions of growth rates when cells are growing in glucose only. The proportion of induced cells is reported for each replicate (Fig. 3C upper panel) while the distribution of growth rates is summarised per condition (Fig. 3C lower panel; mean ± s.d.); the corresponding histograms of growth rates are shown for each replicate in Fig. S13.

We also set out to establish whether the linear dependency of CRP activity to growth rate reported in bulk (often called the C-line) [3] holds at the single-cell level when the growth rate is modulated by the sugar concentration. For this, bacteria were grown in glucose at different concentrations, each supplemented with 200 µM IPTG in order to inhibit LacI repression. Using the filtering criteria described above, we computed growth rate, and the concentration (as the mean of the concentration measured over time) for each cell trace. Importantly, since the growth rate varies over a broad range, the fraction of GFP bleached due to repeated illumination is different between conditions, and the average diameter of the cell also varies. To estimate changes in diameter, we fitted an ellipse to each cell in GFP images for a few randomly selected frames and cells for each condition. Since the estimation of the diameter is noisy relative to its biological fluctuations, we did not intend to measure the diameter on each frame but rather estimated the average diameter in each condition as the average length of the ellipse's small axis. For each observation the cell volume was computed assuming a spherocylinder shape, using this average diameter and the instantaneous length of the cell. To estimate $c$, the concentration of GFP, from $c_u$, the concentration of unbleached GFP, we assumed that the volumic production of GFP $q$ and the growth rate $\lambda$ are approximately constant over the cell cycle. At steady state, $c_u = q/(\lambda + \beta)$ and the concentration of total GFP is $c = q/\lambda$, where $\beta$ is the photobleaching rate. It follows that $c = c_u(1 + \beta/\lambda)$. Thus, for each cell, we estimated the average concentration of all LacZ-GFP molecules (bleached and unbleached) by correcting the average of the measured concentrations, with the photobleaching rate and the average growth rate. We report the joint distributions as the mean ± s.d. of growth rate and concentration for each replicate (Fig. 3B).

**Single-cell level growth and induction** Glucose concentrations at which induced and uninduced cells coexist (32 and 64 µM) display interesting features: the growth rate achieved by induced and uninduced cells (which are expected to feed on lactose and glucose, respectively) is similar (Fig. 3C, Fig. S13), and the fraction of induced cells takes longer to stabilize. Contrary to other conditions with more dissimilar growth rates, we observed that individual cells could switch the expression of the *lac* operon dynamically on and off during the experiments (Fig. 3D). We set out to establish whether those switches could be attributed to GCS at the single-cell level. In this context, the signature of GCS would be that cells switching on the expression of the *lac* operon typically grow slower than average before doing so, and inversely for cells switching off.

In the previous section, we described the analysis of growth and induction at the population level, for which it was sufficient to estimate the growth rate and the induction status for whole cell cycle traces. In order to relate the growth rate to the induction dynamics at the single-cell level, it was necessary to get accurate estimates of the instantaneous growth rate, LacZ-GFP production rate and concentration. To do this, our lab has developed a Bayesian procedure which uses Gaussian process priors for both the volumic production rate and growth rate. The inference software RealTrace (`https://github.com/nimwegenLab/RealTrace`) first optimizes the Gaussian process prior by finding the combination of parameters (average, variance, and correlation times of both growth and volumic production rates, as well as the size of measurement noise in cell size and total fluorescence) that maximizes the likelihood of all the data of an experiment. With this maximum likelihood prior, the program calculates the posterior distributions over cell sizes, total fluorescence, instantaneous growth rate, and instantaneous volumic production rates for each cell at each time point.

We defined induced- and uninduced-states based on the distributions of concentration and volumic production observed in conditions where practically all cells are either induced (lactose 0.02%) or uninduced (lactose 0.02% + glucose 0.02%). On a logarithmic scales, both those variables are well separated by a threshold at 5.5. An on-switch of the *lac* operon expression was defined as going from a volumic production and a concentration below this threshold to both variables above the threshold, and inversely for an off-switch, as illustrated on Fig. 3E. Therefore, a cell whose concentration would cross the production threshold not long enough for the concentration to increase above the threshold would not qualify as switching on, and *vice versa*. In most cases, switches span over multiple cell cycles. We therefore considered switches based on lineages of three cells by appending, to every cell, the data from its parent cell, and average from its daughter cells. The time of the switch was then defined as the time at which the volumic production rate of this lineage crossed the threshold (see time 0 on Fig. S14A). Since the numbers of cells switching the expression of the *lac* operon dynamically is small compared to that of cells being stably induced or uninduced, we have analyzed 45 and 43 supplementary growth channels in the 32 µM and 64 µM conditions, respectively. In total, we could analyze 332 cells switching on and 145 cells switching off in the 32 µM condition (2.8% and 1.2% of the cells), and 142 cells switching on and 81 cells switching off in the 64 µM condition (2.3% and 1.3% of the cells).

Switching lineages were aligned by the time of the switch in order to compute the average growth and volumic production rates as a function of time before and after the switch, using half hours windows around the switch. Comparing the growth rates of cells switching on/off to the average growth rate of induced and uninduced cells in those conditions showed that cells switching on the expression of the *lac* operon typically grew slower than average before doing so, while cells switching off the expression of the *lac* operon typically grew faster than average before doing so (see Fig. S14B). Similarly, comparing the growth rate distributions of cells switching and not switching in the two quadrants of the concentration *vs* volumic production rate space that correspond to transitions states between induced and un-induced, we could show that cells switching off the expression of the *lac* operon typically have a lower growth rate than the other cells that have high concentrations, while cells switching on the expression of the *lac* operon typically have a higher growth rate than the other cells that have a low concentration (see Fig. 3F and Fig. S14B). This difference of growth rate is significant (p-value are indicated in Table S3 for a Welch Two Sample t-test).

Together, those observations demonstrate that GCS impacts induction dynamics at the level of individual cells, and illustrate how this phenomenon shapes *E. coli*'s phenotypic heterogeneity in environments where no strategies is substantially better than the other.

**Table S1.** List of experiments on growth-coupled sensitivity during transient growth arrest (Fig. 2D) with summary statistics.

| condition | date | # growth channels | # cells at switch | # estimated lags | # arrested cells at switch |
|---|---|---|---|---|---|
| to glyc TMG20 | 20180712 | 31 | 174 | NA | 2 |
| to lactulose | 20181024 | 31 | 124 | NA | 3 |
| to lactulose TMG20 | 20181008 | 40 | 147 | 134 | 5 |
| to lactulose TMG20 | 20181009 | 37 | 139 | 124 | 4 |

**Table S2.** List of experiments on the concentration-dependent sugar preference (Fig. 3B-C) with summary statistics. Experiments that have been discarded from further analysis are greyed out.

| treatment | [glucose] ($\mu$M) | date | # analysed cells | prop. induced | # growth channels | # frames |
|---|---|---|---|---|---|---|
| glucose only | 2 | 20210122 | 70 | 0 | 12 | 280 |
| glucose only | 2 | 20210305 | 85 | 0 | 10 | 229 |
| glucose only | 2 | 20210513 | 195 | 0 | 11 | 960 |
| glucose only | 4 | 20210513 | 573 | 0 | 10 | 960 |
| glucose only | 8 | 20210513 | 835 | 0 | 10 | 909 |
| glucose only | 8 | 20210708 | 642 | 0 | 10 | 960 |
| glucose only | 16 | 20210513 | 1383 | 0 | 10 | 960 |
| glucose only | 32 | 20210513 | 629 | 0 | 5 | 960 |
| glucose only | 32 | 20210708 | 1269 | 0 | 10 | 960 |
| glucose only | 64 | 20210513 | 2070 | 0 | 10 | 960 |
| glucose only | 128 | 20210708 | 2115 | 0 | 10 | 960 |
| glucose only | 1009 | 20210708 | 3327 | 0 | 11 | 960 |
| glucose only | 10092 | 20210708 | 3409 | 0 | 10 | 960 |
| with lactose 0.55 mM | 0 | 20210122 | 119 | 1 | 11 | 280 |
| with lactose 0.55 mM | 0 | 20210305 | 657 | 1 | 10 | 370 |
| with lactose 0.55 mM | 2 | 20210122 | 121 | 0.98 | 6 | 251 |
| with lactose 0.55 mM | 2 | 20210305 | 295 | 0.95 | 10 | 308 |
| with lactose 0.55 mM | 4 | 20210122 | 5 | 1 | 11 | 124 |
| with lactose 0.55 mM | 4 | 20210305 | 550 | 0.99 | 10 | 370 |
| with lactose 0.55 mM | 8 | 20210122 | 54 | 1 | 7 | 241 |
| with lactose 0.55 mM | 8 | 20210305 | 404 | 1 | 10 | 273 |
| *with lactose 0.55 mM* | *16* | *20210122* | *47* | *0* | *10* | *260* |
| with lactose 0.55 mM | 16 | 20210305 | 436 | 1 | 10 | 251 |
| with lactose 0.55 mM | 16 | 20210504 | 2022 | 1 | 10 | 642 |
| *with lactose 0.55 mM* | *32* | *20210122* | *166* | *0.42* | *10* | *280* |
| *with lactose 0.55 mM* | *32* | *20210305* | *330* | *0.33* | *10* | *284* |
| with lactose 0.55 mM | 32 | 20210504 | 1371 | 0.92 | 10 | 642 |
| with lactose 0.55 mM | 64 | 20210504 | 829 | 0.44 | 10 | 642 |
| with lactose 0.55 mM | 64 | 20210506 | 209 | 0.88 | 10 | 432 |
| with lactose 0.55 mM | 128 | 20210504 | 1889 | 0 | 10 | 642 |
| with lactose 0.55 mM | 128 | 20210506 | 45 | 0 | 3 | 372 |
| with lactose 0.55 mM | 128 | 20210708 | 3186 | 0.03 | 11 | 960 |
| with lactose 0.55 mM | 1009 | 20210122 | 378 | 0 | 10 | 280 |
| with lactose 0.55 mM | 1009 | 20210305 | 942 | 0 | 10 | 370 |
| with IPTG 200$\mu$M | 2 | 20210504 | 257 | 1 | 11 | 642 |
| with IPTG 200$\mu$M | 2 | 20210506 | 62 | 0.98 | 11 | 432 |
| with IPTG 200$\mu$M | 2 | 20210513 | 263 | 1 | 11 | 960 |
| with IPTG 200$\mu$M | 8 | 20210504 | 513 | 1 | 10 | 642 |
| with IPTG 200$\mu$M | 8 | 20210506 | 114 | 1 | 8 | 432 |
| with IPTG 200$\mu$M | 32 | 20210504 | 1260 | 1 | 10 | 642 |
| with IPTG 200$\mu$M | 32 | 20210506 | 645 | 1 | 10 | 432 |
| with IPTG 200$\mu$M | 128 | 20210506 | 709 | 1 | 17 | 432 |
| with IPTG 200$\mu$M | 128 | 20210513 | 2541 | 1 | 11 | 960 |
| M9 zero | 0 | 20210708 | 73 | 0 | 10 | 960 |

| Condition | Parameter space | p-value |
|---|---|---|
| Lac+glu 32 µM | Low [GFP], high production | $1.98 \times 10^{-8}$ |
| Lac+glu 32 µM | High [GFP], low production | $1.58 \times 10^{-7}$ |
| Lac+glu 64 µM | Low [GFP], high production | $9.15 \times 10^{-5}$ |
| Lac+glu 64 µM | High [GFP], low production | $1.45 \times 10^{-2}$ |

**Table S3.** Significance of the differences of mean growth rate between cells switching and cells not switching in the different conditions and different sectors of the parameter space, calculated with a Welch two sample t-test.

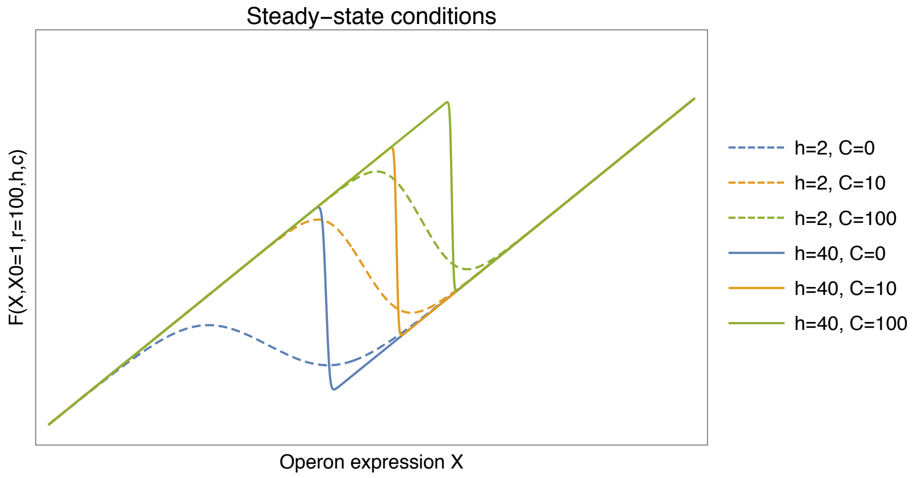


**Figure S1.** Steady-state conditions for the auto-activating two-component system coupled to an external signal $s$. Shown is the function $F(X/X_0, r, h, C)$ of equation (S21) as a function of the protein concentration $X$ for a critical level $X_0 = 1$, fold-change $r = 100$, and Hill coefficients $h = 2$ (dashed) or a high value $h = 40$ (solid), and for $C = 0$ (blue), $C = 10$ (orange), and $C = 100$ (green).

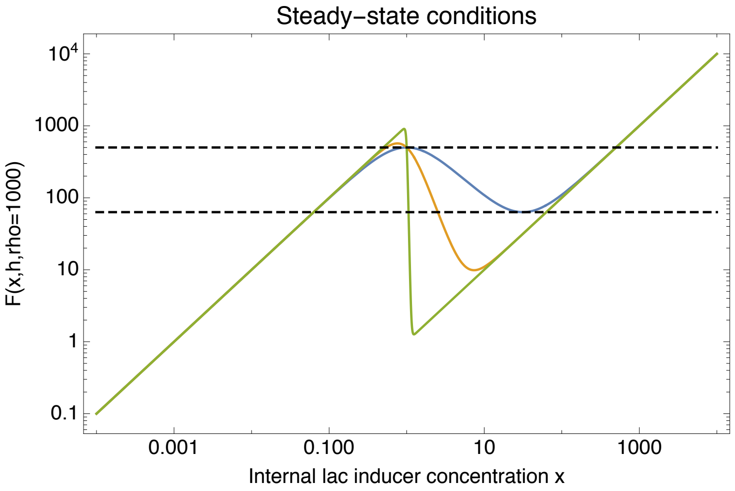


**Figure S2.** Illustration of the steady-state condition for the *lac* operon model (Eq. S33). The function $F(x, h, \rho)$ is shown as a function of the intracellular inducer concentration $x$ (horizontal axis) for Hill coefficients $h = 2$ (blue), $h = 4$ (orange) and $h = 50$ (green). The dashed black lines show the critical values of $F$ that define the bistable regime for $h = 2$.

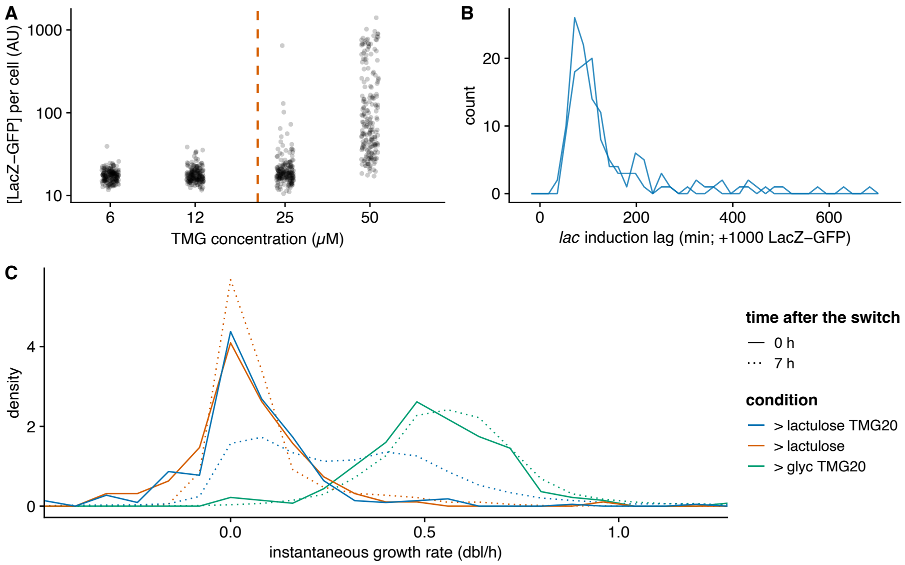


**Figure S3.** **A**. Characterization of the lower threshold for the *lac* operon bistability under TMG induction for strain ASC662. For each TMG concentration, 400 cells were measured; the orange dashed line indicates the concentration used for experiments with lactulose (20 µM; Fig. 2D). **B**. Distribution of the lag times to induction of the *lac* operon exposed to 0.2% lactulose supplemented with 20 µM TMG (2 independent replicates). **C**. Distributions of instantaneous growth rates for bacteria after the switch to media with 20 µM TMG in different nutrients, measured over 1 h right after the switch or after 7h of induction. Growth on glycerol isn't affected by adding TMG as shown by the fact that growth rates are the same right after the switch and 7h later (green; more than 200 cells per histogram); to the contrary, all cells have stopped growing immediately after the switch to lactulose (orange and blue solid lines) and growth resumes only for a subset of cells in the presence of TMG (blue dotted line; ≈ 450 cells).

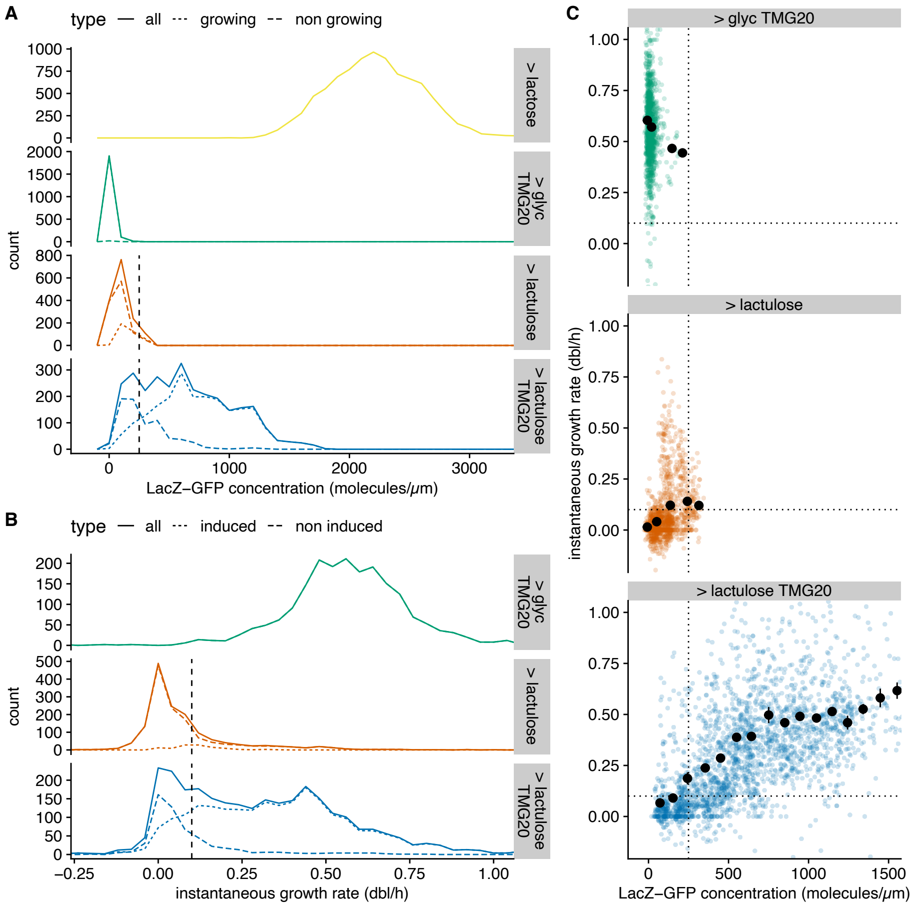


**Figure S4.** **A-B**. Distributions of single-cell LacZ-GFP levels (A) and instantaneous growth rates (B) for bacteria 7 h after a switch from glycerol minimal media to either minimal media supplemented with either lactose (yellow), glycerol with 20 µM of TMG (green), lactulose (orange), or lactulose with 20 µM of TMG (blue). While no *lac* induction occurs with 20 µM TMG in glycerol or lactulose only, the majority of cells express LacZ-GFP when growing in lactulose supplemented with the same concentration of TMG. Note that, since most cells tend to express a very low amount of LacZ-GFP even in the absence of TMG (orange), we set a threshold for expression attributable to TMG at 250 molecules/µm (dashed line in A). Similarly, since very slow growth is observed on lactulose without TMG, we set a threshold to distinguish growing cells at 0.1 dbl/h (dashed line in B). Notably, growth rate and LacZ-GFP levels are correlated in lactulose supplemented with TMG (see panel C) and the distribution of LacZ-GFP levels can be decomposed into two well-separated underlying distributions simply by distinguishing growing from non-growing cells, and *vice versa*. Lastly, comparing the distributions of LacZ-GFP levels in 0.2% lactulose supplemented with 20 µM TMG (blue) and in 0.2% lactose (yellow) clearly shows that this low TMG concentration does not allow full induction of the *lac* operon. **C**. Single-cell instantaneous growth rates versus LacZ-GFP levels for bacteria 7h after a switch to glycerol with 20 µM of TMG (green), lactulose (orange), and lactulose with 20 µM of TMG (blue). The clear correlation between growth rate and LacZ-GFP level in lactulose supplemented with TMG indicates that *lac* expression is limiting growth in this condition as expected at such a low concentration of inducer [19]. Dotted lines show the threshold for growth and induction in panels A and B. In all panels, all growth rates and LacZ-GFP levels are estimated over 1 h of measurements, i.e. between 7 and 8 h after the switch.

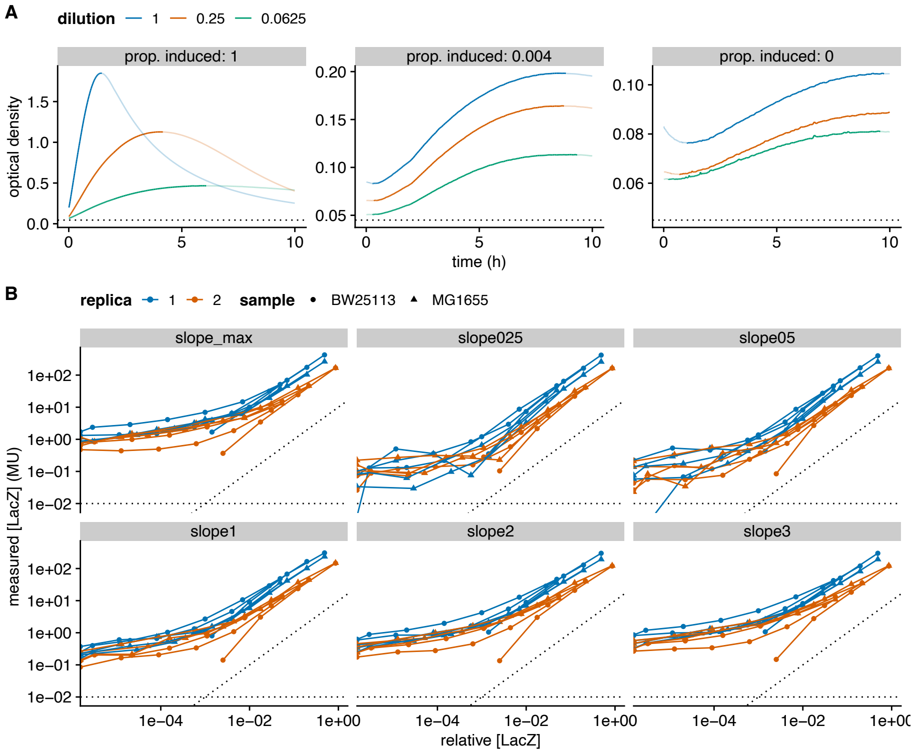


**Figure S5.** Calibration of the Miller assay. **A**. Example time series of $OD_{420}$ with continuous recording; the concentration of β-galactosidase (LacZ) is varied by changing both the ratio of induced and uninduced cultures in the sample (panels), and the total cell concentration (colours indicating the dilution relative to a sample where $OD_{600} = 0.8$). Note that the kinetics is not monotonic, and that $OD_{420}$ starts by decreasing at low LacZ and bacteria concentration. To estimate the β-galactosidase concentration, we focus on the first monotonically increasing time window (opaque lines). **B**. Comparison of several variables to estimate β-galactosidase (LacZ) concentration from the $OD_{420}$ kinetics. In each panel, the measured LacZ concentration is plotted for each sample (vertical axis) against the relative LacZ concentration (derived from the proportion of induced culture and the cell concentration; horizontal axis); this relationship is expected to be linear (dotted lines are a guide to the eye with slopes 0 and 1). Panel titles indicate the duration in hours used to estimate LacZ concentration (*slope025*: 15 min, *slope05*: 30 min, *slope1*: 1 h, etc); colours indicate biological replicates, and each line corresponds to a technical replicate.

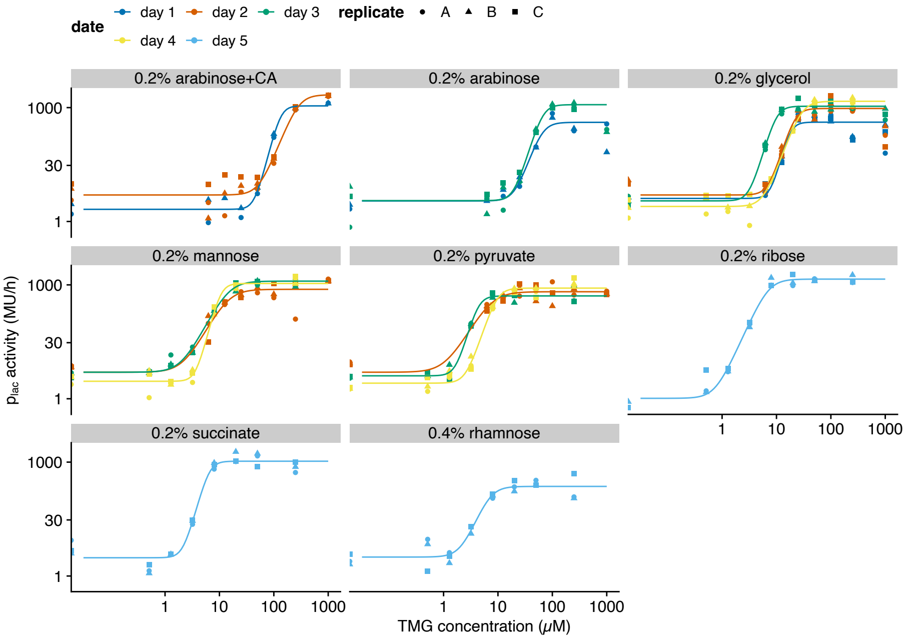


**Figure S6.** Induction curves of the *lac* operon with TMG for several nutrients, measured with Miller assay (as in Fig. 2E, here with all conditions and replicates). Lines show fits to a Hill function for each biological replicate, with (up to) three data points at a given concentration corresponding to three technical replicates (same colour).

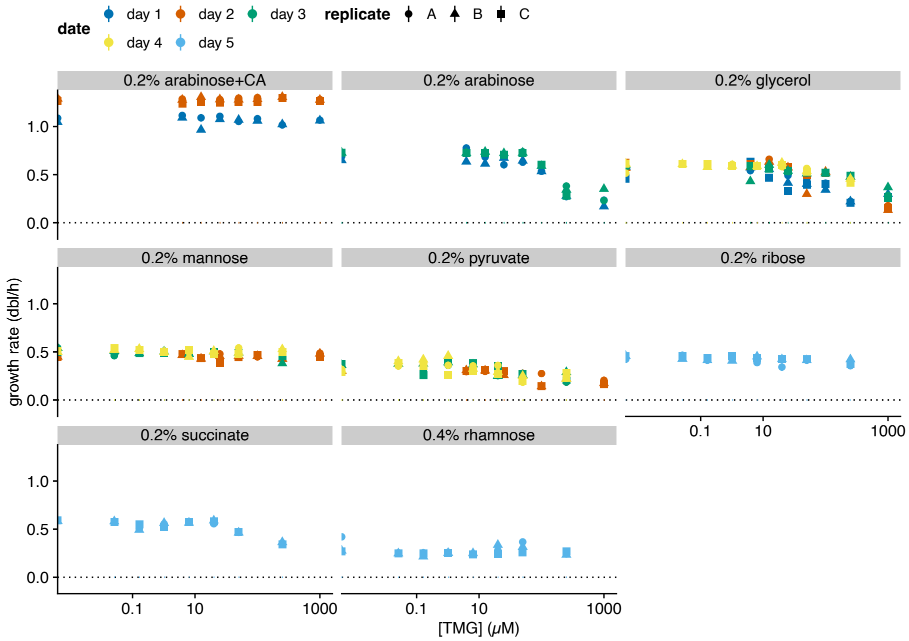


**Figure S7.** Growth rate as a function of TMG concentration, stratified by nutrient, for experiments in balanced exponential growth where the growth rate is modulated by changing the nutrients (Fig. 2E-F). Error bars show standard error of the slope of the linear fit of $\log(OD_{600})$ versus time.

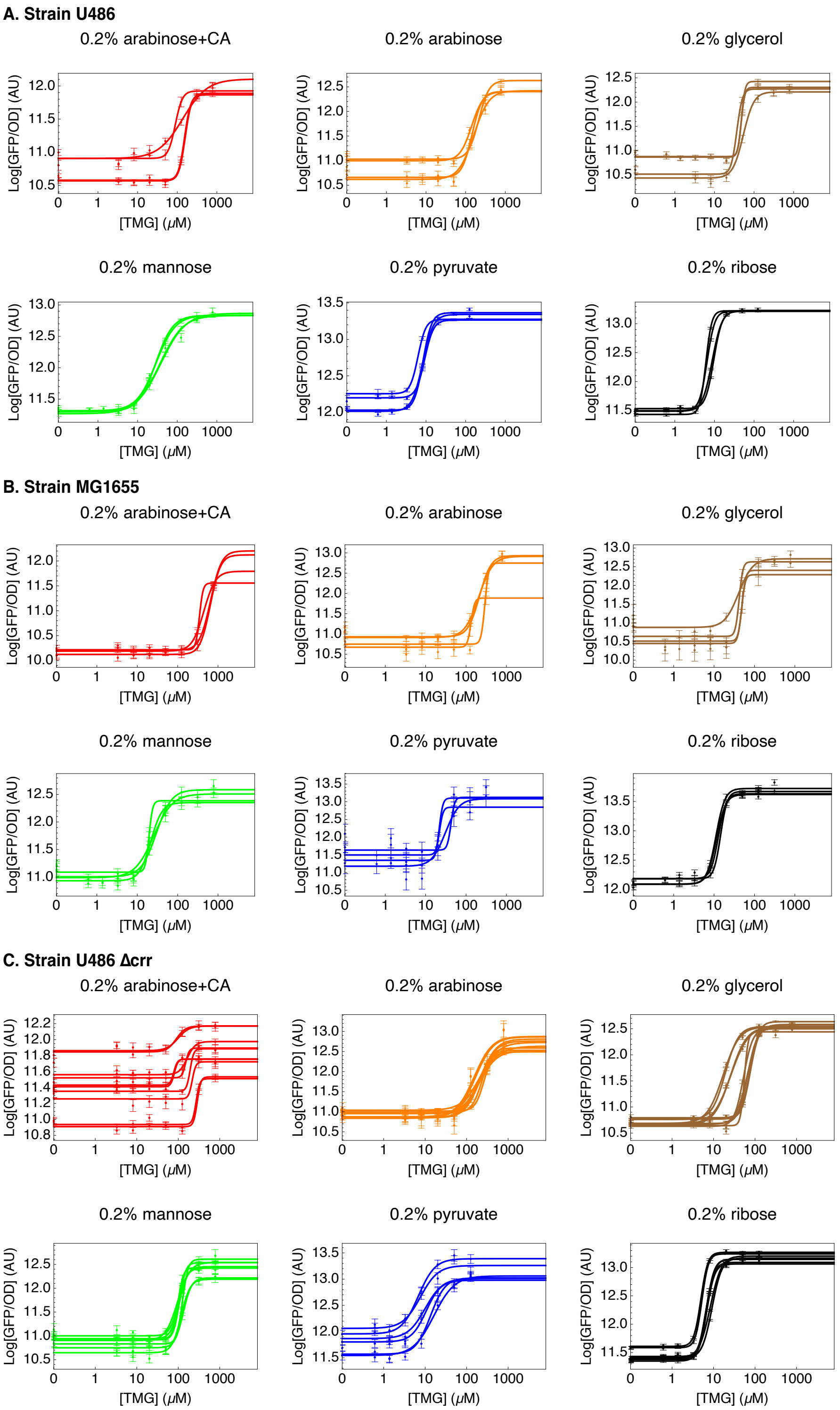


**Figure S8.** Induction curves of the *lac* operon with TMG for several nutrients, measured with LacZ-GFP fluorescence for three different strains. Lines show fits to a Hill function for each biological replicate. **A**. Induction curves for strain U486 (MG1655 ΔcpdA ΔcyaA) supplemented with 1mM cAMP. **B**. Induction curves for the wild-type strain MG1655. **C**. Induction curves for strain U486 Δcrr, supplemented with 1mM cAMP, where the inducer exclusion effect is abolished.

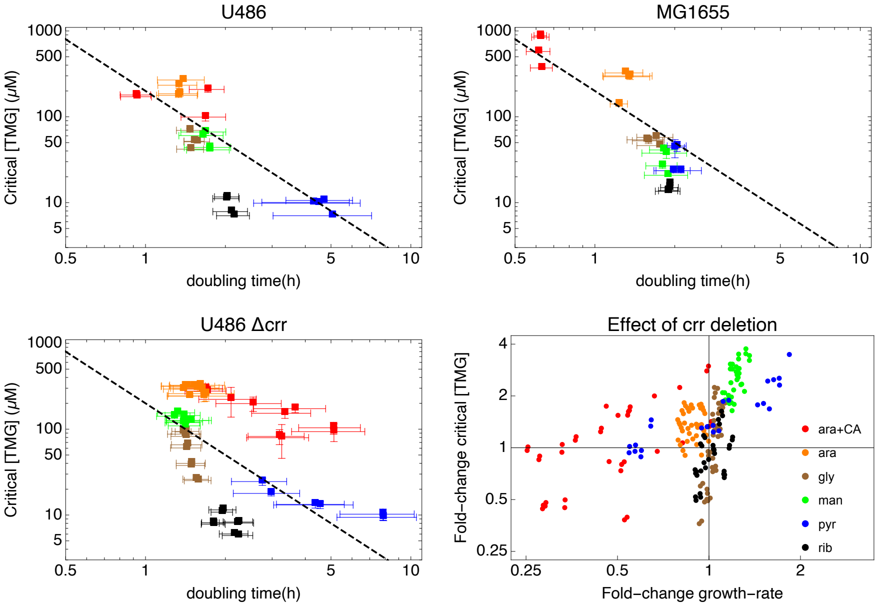


**Figure S9.** Growth-coupled sensitivity plots obtained for three different strains, when the growth rate is modulated with nutrients. The critical concentration fitted in Fig. S8 is plotted against the growth rate measured for the same replicate. The bottom right panel shows fold-change of the critical TMG concentration vs the fold change of growth rate between U486 Δcrr and U486; this highlights that inducer exclusion is not underlying the changes of critical concentration observed when the growth rate is modulated with different nutrients.

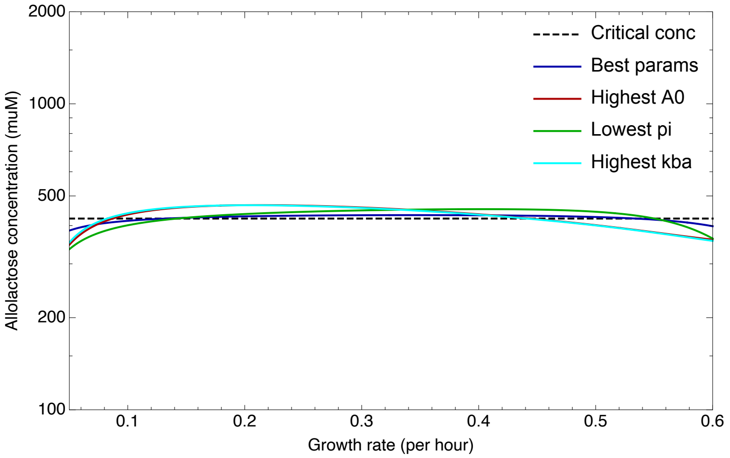


**Figure S10.** Example parameter regimes of LacZ metabolism supporting optimal carbon source switching. In order for the critical external lactose concentration to track the inverse Monod equation for lactose, the steady-state internal allolactose concentration at the critical LacY and LacZ levels should equal $x_{\text{crit}} \approx 420\mu M$ independent of growth rate (dotted black line, see SI section 4). Colored curves show steady-state intracellular allolactose concentrations as a function of growth rate for example kinetic LacZ parameter settings that approximate this behavior:

- Best parameter settings
  $A_0 = 29\mu M, p_i = 910, k_{bl} = 73.5(\mu M h)^{-1}, k_{ba} = 0.05(\mu M h)^{-1}, b_r = 0.79$ (blue),
- High $A_0$ parameters
  $A_0 = 128\mu M, p_i = 758, k_{bl} = 5.7(\mu M h)^{-1}, k_{ba} = 0.05(\mu M h)^{-1}, b_r = 0.27$ (red),
- Low $p_i$ parameter settings
  $A_0 = 75\mu M, p_i = 212, k_{bl} = 42.5(\mu M h)^{-1}, k_{ba} = 0.05(\mu M h)^{-1}, b_r = 1.37$ (green),
- High $k_{ba}$ parameter settings
  $A_0 = 1.32\mu M, p_i = 632, k_{bl} = 455(\mu M h)^{-1}, k_{ba} = 3.31(\mu M h)^{-1}, b_r = 0.46$ (cyan).

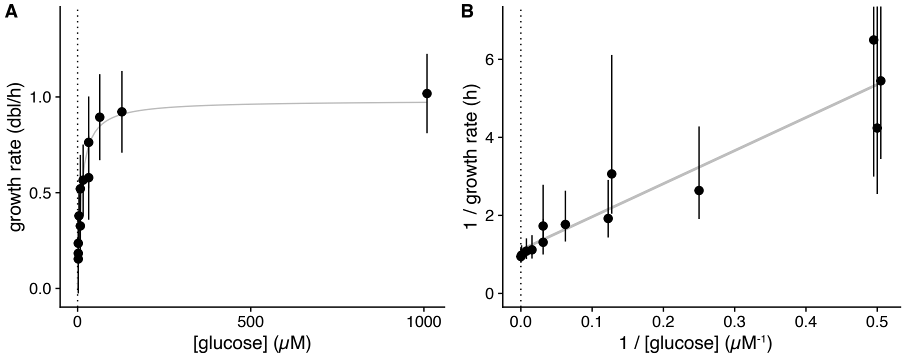


**Figure S11.** Distributions of single-cell growth rates for bacteria grown at different glucose concentrations. **A**. Distribution of single-cell growth rates (mean ± standard-deviation) as a function of glucose concentration. The grey curve shows a least-squares fit to a hyperbolic function (i.e. Monod curve; $K_m = 10 \pm 2$ µM and $\lambda_{max} = 0.98 \pm 0.05$ dbl/h). **B**. The same distributions as shown in panel A, but now showing the inverse of growth rate as a function of the inverse of glucose concentration, which should show a linear dependence if the growth rates follow a hyperbolic Monod function. The grey line is a linear fit, which confirms that single-cell growth rates at low glucose concentrations indeed follow a Monod function.

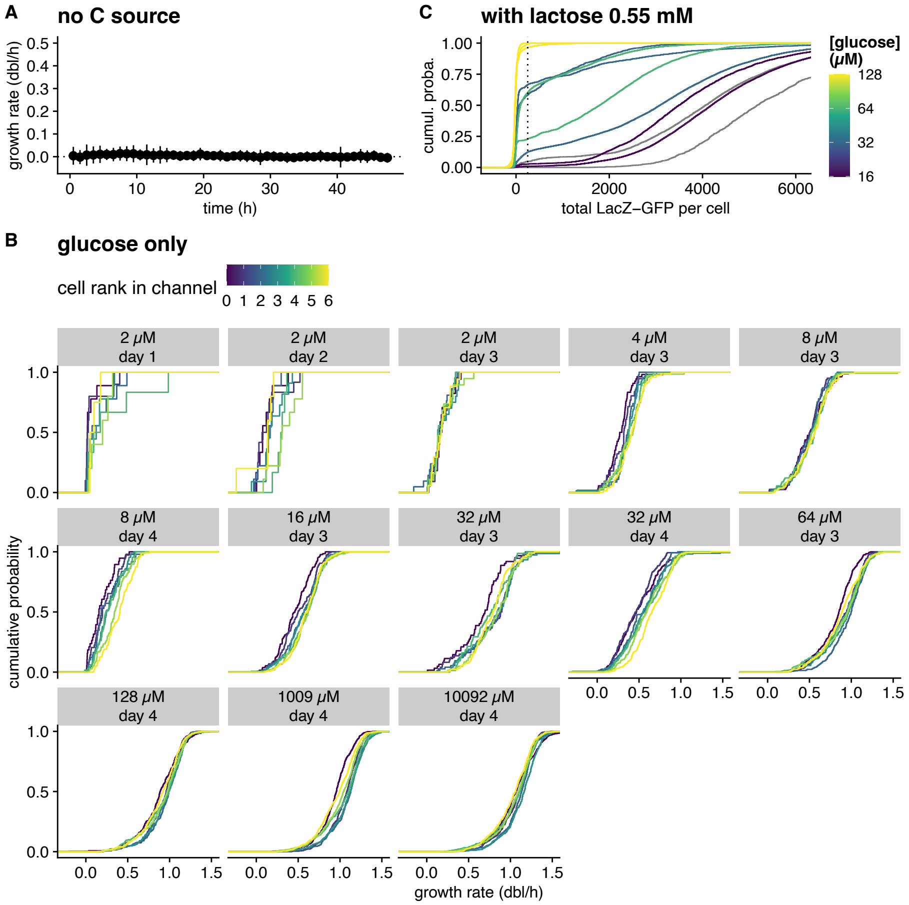


**Figure S12.** Controls for the glucose-lactose concentration-dependent preference (shown in Fig. 3B&C). **A**. Growth rate as a function of time for bacteria exposed to M9 without a carbon source. The growth rate is measured for each cell as the slope of the linear regression of log(length) as a function of time, within time windows of 1 h (dots and errors bars show the mean and standard deviation over 72 cells). Since the growth rate is null during this entire experiment, we conclude that growth observed in the experiments with very low glucose concentration does not come from residual nutrients present in the setup (reservoirs, tubing, etc). **B**. Cumulative distributions of growth rates stratified by cell position in the GC (in colour; rank 0 corresponds to the mother cell), for each replicate where bacteria were exposed to glucose only. The lack of systematic dependence of the growth rate on the cell's position in the channel supports that there are no gradients of nutrients within the GCs. **C**. Cumulative distributions of all measurements of total LacZ-GFP intensity per cell, at several glucose concentrations in experiments with a mixture of glucose and lactose. The distribution corresponding to lactose (0.2%) without glucose is shown in gray, while the distribution corresponding to abundant glucose (128 µM) in the mixture is shown in yellow. At all intermediate concentrations of glucose, the distributions are bimodal, and the mode of the uninduced subpopulations (corresponding to the steep increase at the left of the cumulative density functions) is well captured using a threshold of 250 LacZ-GFP per cell (vertical dotted line). This induction threshold is used to compare the growth rate of induced and uninduced bacteria in Fig. 3C and in Fig. S13.

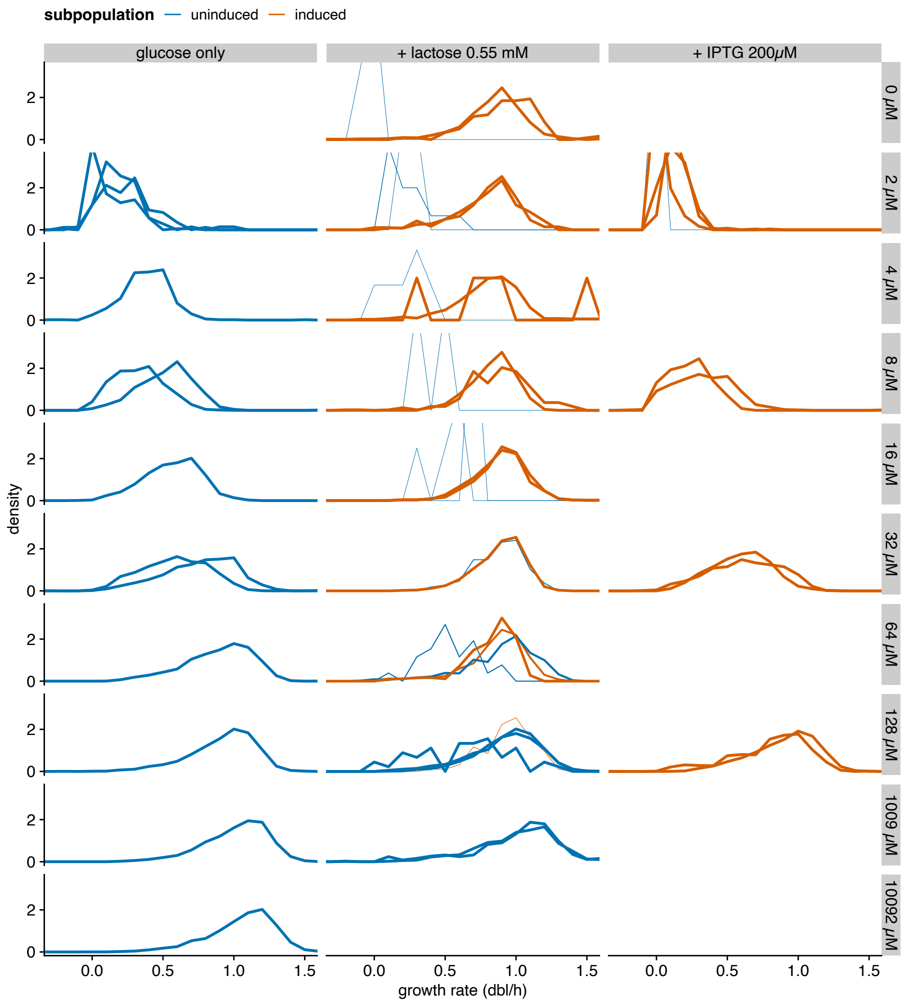


**Figure S13.** Distributions of growth rate for single cells grown on glucose at different concentrations (left), supplemented with 0.55mM (i.e. 0.02%) lactose (middle) or 200 µM IPTG (right). Each replicate experiment is shown independently (more than 70 cells in each; median ≈ 600 cells); cells are stratified into induced or uninduced based on their absolute fluorescence level (> 250 GFP molecules). The thickness of the lines indicates the proportion of the corresponding category in the replicate.

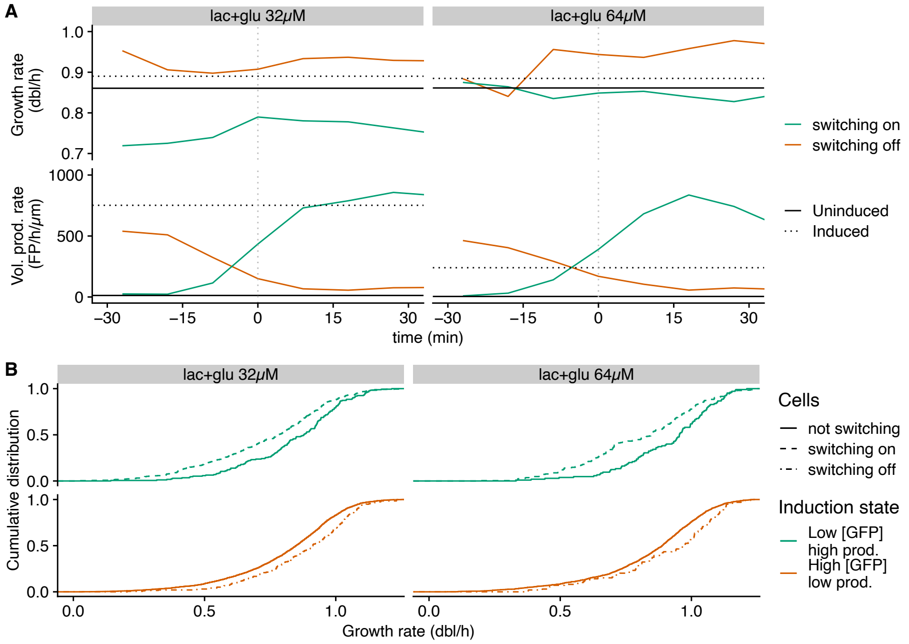


**Figure S14.** Growth coupled sensitivity impacts the switching dynamics of the *lac* operon at the single-cell level, in mixtures of glucose and lactose where both nutrients support comparable growth (0.55 mM lactose + 32 or 64 μM glucose). **A**. Medians of growth and volumic-production rates across time for lineages switching on/off (green/orange, respectively), compared to induced/uninduced cells in the same conditions (solid/dotted horizontal lines, respectively). Time 0 is defined as the moment where the cell crosses the production threshold (vertical dotted line). **B**. Cumulative distributions of growth rate for cells in different sectors of the parametric space defined in Fig. 3E (colors), stratified by the switch of the *lac* operon (shown as different types of dashed lines).

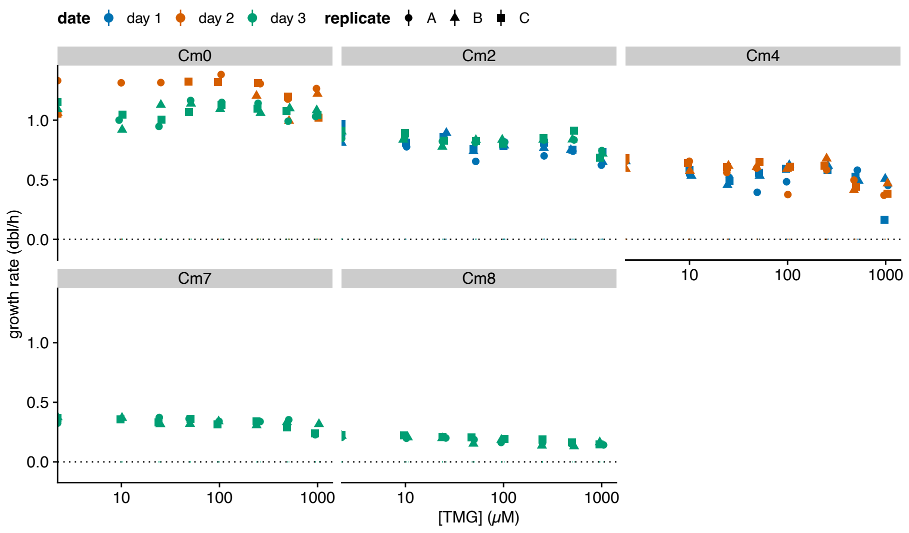


**Figure S15.** Growth rate as a function of TMG concentration, stratified by Cam concentration, for experiments in balanced exponential growth where the growth rate is modulated by subinhibitory Cam level (Fig. 3G-H). Error bars show standard-errors of the slope of the linear fit of log($OD_{600}$) versus time.

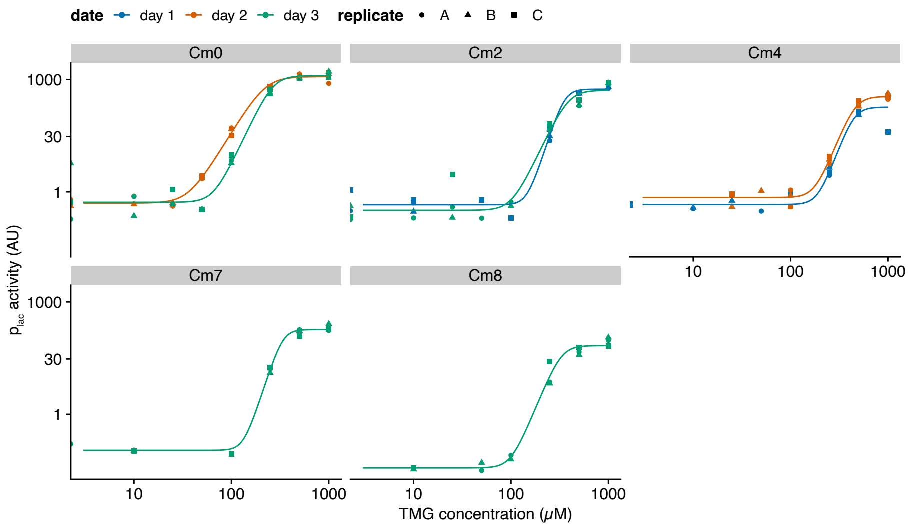


**Figure S16.** Induction curves of the *lac* operon with TMG for increasing subinhibitory Cam levels, measured with Miller assay. Lines show fits to a Hill function, with (up to) three data points at a given concentration corresponding to three technical replicates (same colour), used to estimate the critical TMG concentration at each Cam concentration (shown in Fig. 3H).

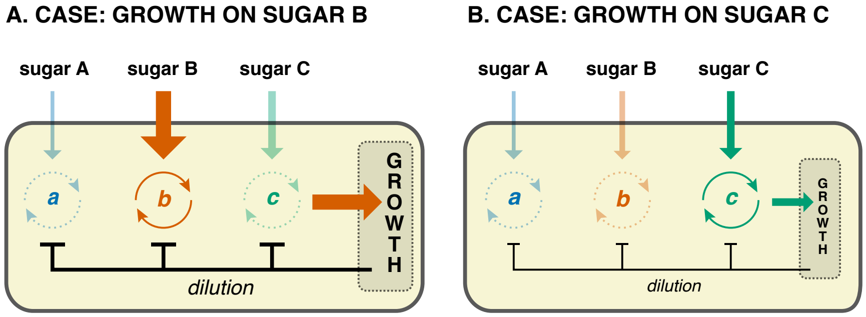


**Figure S17.** Schematic of the circuitry through which GCS can implement optimal concentration-dependent sugar preferences. **A**. The cartoon depicts a bacterial cell that contains three regulatory positive feedback switches (a, b, and c) for the catabolic genes of sugars A, B, and C. The induction of each of the regulatory switches is positively affected by the influx of its corresponding sugar (vertical arrows) and negatively affected by dilution (black inhibitory arrows). The strength of this dilution is determined by the growth rate of the cell, which in turn is set by the flux of the sugar whose switch is induced (switch b and red horizontal arrow). Consequently, only the switch corresponding to the sugar with the highest flux will be induced. For all others the negative effect of dilution is larger than the positive effect of the influx of their sugar. **B**. The identical cell in another environment where the flux of sugar B is reduced. In this environment, switch C will be induced even though its flux is the same as in panel A.
